# Non-lytic spread of poliovirus requires the nonstructural protein 3CD

**DOI:** 10.1101/2024.10.18.619132

**Authors:** David Aponte-Diaz, Jayden M. Harris, Tongjia Ella Kang, Victoria Korboukh, Mohamad S. Sotoudegan, Jennifer L. Gray, Neela H. Yennawar, Ibrahim M. Moustafa, Andrew Macadam, Craig E. Cameron

## Abstract

Non-enveloped viruses like poliovirus (PV) have evolved the capacity to spread by non-lytic mechanisms. For PV, this mechanism exploits the host secretory autophagy pathway. Virions are selectively incorporated into autophagosomes, double-membrane vesicles that travel to the plasma membrane, fuse, and release single-membrane vesicles containing virions. Loading of cellular cargo into autophagosomes relies on direct or indirect interactions with microtubule-associated protein 1B-light chain 3 (LC3) that are mediated by motifs referred to as LC3-interaction regions (LIRs). We have identified a PV mutant with a severe defect in non-lytic spread. An F-to-Y substitution in a putative LIR of the nonstructural protein 3CD prevented virion incorporation into LC3-positive autophagosomes and virion trafficking to the plasma membrane for release. Using high-angle annular dark-field scanning transmission electron microscopy to monitor PV-induced autophagosome biogenesis, for the first time, we show that virus-induced autophagic signals yield normal autophagosomes, even in the absence of virions. The F-to-Y derivative of PV 3CD was unable to support normal autophagosome biogenesis. Together, these studies make a compelling case for a direct role of a viral nonstructural protein in the formation and loading of the vesicular carriers used for non-lytic spread that may depend on the proper structure, accessibility, and/or dynamics of its LIR. The studies of PV 3CD protein reported here will hopefully provoke a more deliberate look at the presence and function of LIR motifs in viral proteins of viruses known to use autophagy as the basis for non-lytic spread.

**IMPORTANCE:** PV and other enteroviruses hijack the cellular secretory autophagy pathway for non-lytic virus transmission. While much is known about the cellular factors required for non-lytic transmission, much less is known about viral factors contributing to transmission. We have discovered a PV nonstructural protein required for multiple steps of the pathway leading to vesicle-enclosed virions. This discovery should facilitate identification of the specific steps of the cellular secretory autophagy pathway and corresponding factors commandeered by the virus and may uncover novel targets for antiviral therapy.

## INTRODUCTION

Poliovirus (PV), the prototypical member of the *Enterovirus* genus of the *Picornaviridae* family of positive-strand RNA viruses, is among the best-understood viruses (1). While PV has been essentially eliminated from developing countries due to effective vaccination measures, global eradication has yet to be certified (2–5). We have used PV as a model system to understand the enzymology and cell biology of viral genome replication (6–9) because of the more than 50 years of work by dozens of investigators establishing a solid foundation of principles governing PV multiplication (10–13).

The latest emerging principle is that non-enveloped picornaviruses spread preferentially by concealing virions within vesicles and using a non-lytic mechanism instead of a lytic mechanism (14–16). Secretory autophagy is the predominant mechanism for the non-lytic spread of PV and other enteroviruses (17–21). The literature supporting this conclusion has interrogated the extent to which what is known about the cellular mechanism of secretory autophagy and corresponding factors align with the virus-induced pathway (22–25). Very little is known about direct contributions of viral factors to non-lytic spread.

We have identified a derivative of the PV nonstructural protein 3CD that causes a defect in the non-lytic spread of the virus. Using a variety of approaches, including high-angle annular dark-field (HAADF) scanning transmission electron microscopy (STEM), we show that 3CD is required for particle movement from the site of assembly into autophagosomes, proper formation of autophagosomes, and movement of virion-containing autophagosomes from the perinuclear region of the cell to the periphery and beyond. We suggest that one or more LC3-interacting regions of 3CD contribute to its post-genome-replication functions and that the multifunctional properties of 3CD are bestowed by its highly tunable, extraordinary conformational dynamics (26–28).

## RESULTS

### A post-genome-replication function for PV 3CD protein

The processivity of nucleotides incorporated per binding event by poliovirus RNA-dependent RNA polymerase (PV RdRp), without any accessory factors, is predicted to approach 10^6^ (29). The best empirical evidence shows that PV RdRp can replicate through 3000 bp of dsRNA without dissociating (29, 30). The processivity of reverse transcriptases does not approach these values (31, 32). Accessory factors enabling processivity are generally required for DNA polymerases to exhibit high processivity (33–35). The structural basis for PV RdRp processivity is not known. One hypothesis has been that the intimate interaction between the fingertips and thumb subdomains of the RdRp yields a completely encircled active site that will not readily dissociate from the RNA template once engaged (**Fig. 1A**).

**Figure 1.**
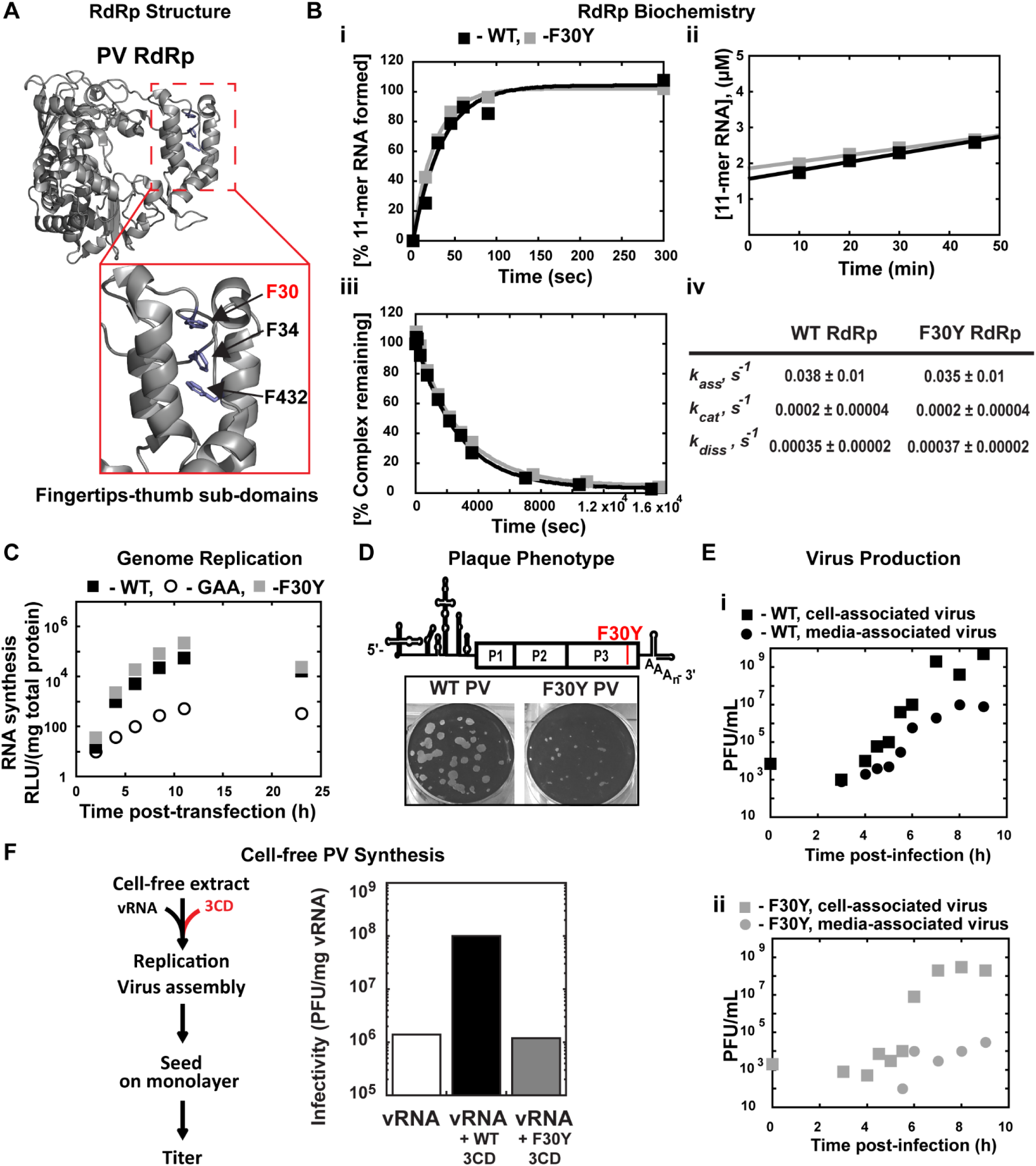
A post-genome-replication function for PV 3CD protein. **(A) PV 3D RdRp structure.** The PV 3D RdRp structure is depicted as a gray ribbon; the structure adopts a canonical right-hand shape with fingers, palm, and thumb subdomains. The red box inset shows a close-up view of the fingertips-thumb subdomain interaction. Residues F30, F34 (fingertips), and F432 (thumb) are highlighted in blue to show the phenylalanine “stacking” interaction that occurs between the fingertips and thumb subdomains. The image was created using the WebLab Viewer (Molecular Simulations Ins., San Diego, CA) program (PDB access code 1RA6). **(B) PV 3D F30Y biochemical properties. (i) Complex assembly kinetics.** Shown are the kinetics of RNA product formation over time. Solid lines represent the best fit of the data to a single exponential with assembly rates (*k_ass_*) of 0.038 ± 0.01 s-1 (WT) and 0.035 ± 0.01 s-1 (F30Y). **(ii) Active site titration.** Shown are the kinetics of RNA product formation over time. The data fit best to a straight line with y-intercepts representing concentrations of the active enzyme with 1.6 μM for WT and 1.8 μM for F30Y, corresponding to 80 and 90% of the total enzyme being “active,” respectively. The steady-state rate of AMP incorporation (*k_cat_*) was 0.0002 ± 0.00004 s-1 for both WT and F30Y. **(iii) Complex dissociation kinetics.** Shown are the kinetics of RdRp-primed-template complex dissociation over time. The solid lines represent the best fit of the data to a single exponential with dissociation rates (*k_diss_*) of 0.00035 ± 0.00002 s-1 (WT) and 0.00037 ± 0.00002 s-1 (F30Y). **(iv) WT and F30Y PV RdRp kinetic parameters.** Table summarizing the kinetic parameters for WT and F30Y PV RdRp. **(C) Genome replication.** Sub-genomic replicon luciferase assay comparing WT and F30Y. Luciferase is measured as a surrogate for genome replication using a relative light unit (RLU) normalized to protein content (µg) from an absorbance measure of the collected lysates at the shown time points. In this assay, an inactive polymerase variant GAA PV controlled for translation and RNA stability during inhibited RNA synthesis. (**D**) **Plaque Phenotype.** A schematic PV genome schematic is shown, highlighting the F30Y mutation placement. Comparison of 50 PFU of WT and F30Y PV. The number of PFUs observed for WT and F30Y PV was essentially the same. However, the F30Y virus produced plaques of smaller size. **(E) Virus Production.** One-step growth curve comparing media-associated (supernatant) and cell-associated (cells) virus collected from WT and F30Y PV infections. Titers were quantified by plaque assay. **(i)** WT PV virus titers shown. **(ii)** F30Y PV virus titers shown. **(F) Cell-free PV Synthesis.** Schematic depicting the cell-free extract assay used to detect assembly stimulation in the context of exogenous viral protein supplementation. The graph shows the cell-free synthesis of PV in the presence of WT and F30Y purified 3CD. Titers were quantified by plaque assay and normalized to the amount of vRNA.

The interface between the fingertips and thumb subdomains is comprised, in part, of three phenylalanine residues: F30 and F34 from the fingertips and F432 from the thumb (**Fig. 1A**). We constructed a PV RdRp derivative in which F30 was changed to Y, this derivative is referred to herein as F30Y. Our rationale was that burying the tyrosine hydroxyl and any associated water molecules would destabilize the interface, creating a derivative with only reduced processivity.

We reasoned that an RdRp derivative exhibiting a destabilized interaction with primed-template would not assemble on the primed-template as well as WT RdRp does. Once assembled, the complex would also be predicted to be less stable, causing an increase in the steady-state rate constant for nucleotide incorporation and rate constant for dissociation of the derivative from the primed-template when compared to WT RdRp.

Interestingly, the biochemical properties of F30Y RdRp were identical to WT (**Fig. 1B**). Because our in vitro tests could have been masking a phenotype that would be observed in cells, we engineered the F30Y derivative into a subgenomic replicon to indirectly monitor genome replication after transfection by measuring luciferase activity. This assay also failed to reveal a phenotype for the F30Y RdRp (**Fig. 1C**). We used a replicon expressing an inactive RdRp (GAA) as a negative control. Luciferase activity produced in this case reflects translation of the transfected RNA without genome replication.

For completeness, we engineered the F30Y change into the viral genome. We did not expect a difference between F30Y PV and WT, but F30Y PV exhibited a small-plaque phenotype when compared to WT (**Fig. 1D**). This phenotype could reflect a reduction in infectious virus produced and/or a reduction in the efficiency of virus spread. We performed a one-step-growth analysis and monitored the production of infectious virus within cells (cell-associated) and the efficiency with which virus was released from cells (media-associated). WT PV reproducibly produced virus on the order of 10^9^ plaque-forming units (pfu) per mL, with 10^7^ pfu/mL detectable in media by the end of the experiment (panel i, **Fig. 1E**). In contrast, F30Y PV exhibited a one-log reduction in overall yield of cell-associated virus and three- to four-log reduction in media-associated virus (panel ii, **Fig. 1E**).

The primary form of the RdRp domain in PV-infected cells is the precursor 3CD (36, 37). Previous studies have implicated this protein as a critical factor for genome replication (38–41) and in steps preceding genome replication, including activation of phospholipid biosynthesis and membrane biogenesis (8, 9). The studies reported above are consistent with 3CD as the mediator of the observed effect and a post-genome-replication function for this protein. It is known that purified 3CD stimulates cell-free synthesis of PV (42) through unknown mechanisms. We expressed and purified the corresponding 3CD derivative (F213Y) but will refer to it as F30Y 3CD to avoid confusion. The F30Y 3CD derivative could not support cell-free synthesis stimulation (**Fig. 1F**), which is consistent with 3CD as the mediator of the observed biological defect and further supports a role for 3CD after genome replication.

### PV 3CD contributes to virion morphogenesis and non-lytic spread

Studies of the enterovirus lifecycle can be synchronized using two inhibitors. First, guanidine hydrochloride (GuHCl) at 2 mM is sufficient to inhibit genome replication by inactivating the ATPase activity of the viral 2C protein (43–45). However, the pioneer round of translation occurs, as the activity of genome-encoded reporters like luciferase can be detected (WT+GuHCl in **Fig. 2A**). Second, 5-(3,4-dichlorophenyl) methyl hydantoin, referred to herein as hydantoin (H) also targets the viral 2C protein, but impairs virion morphogenesis without any impact on genome replication (WT+H in **Fig. 2A**) (46, 47), although at concentrations higher than 50 µg/mL, hydantoin can inhibit genome replication (48).

**Figure 2.**
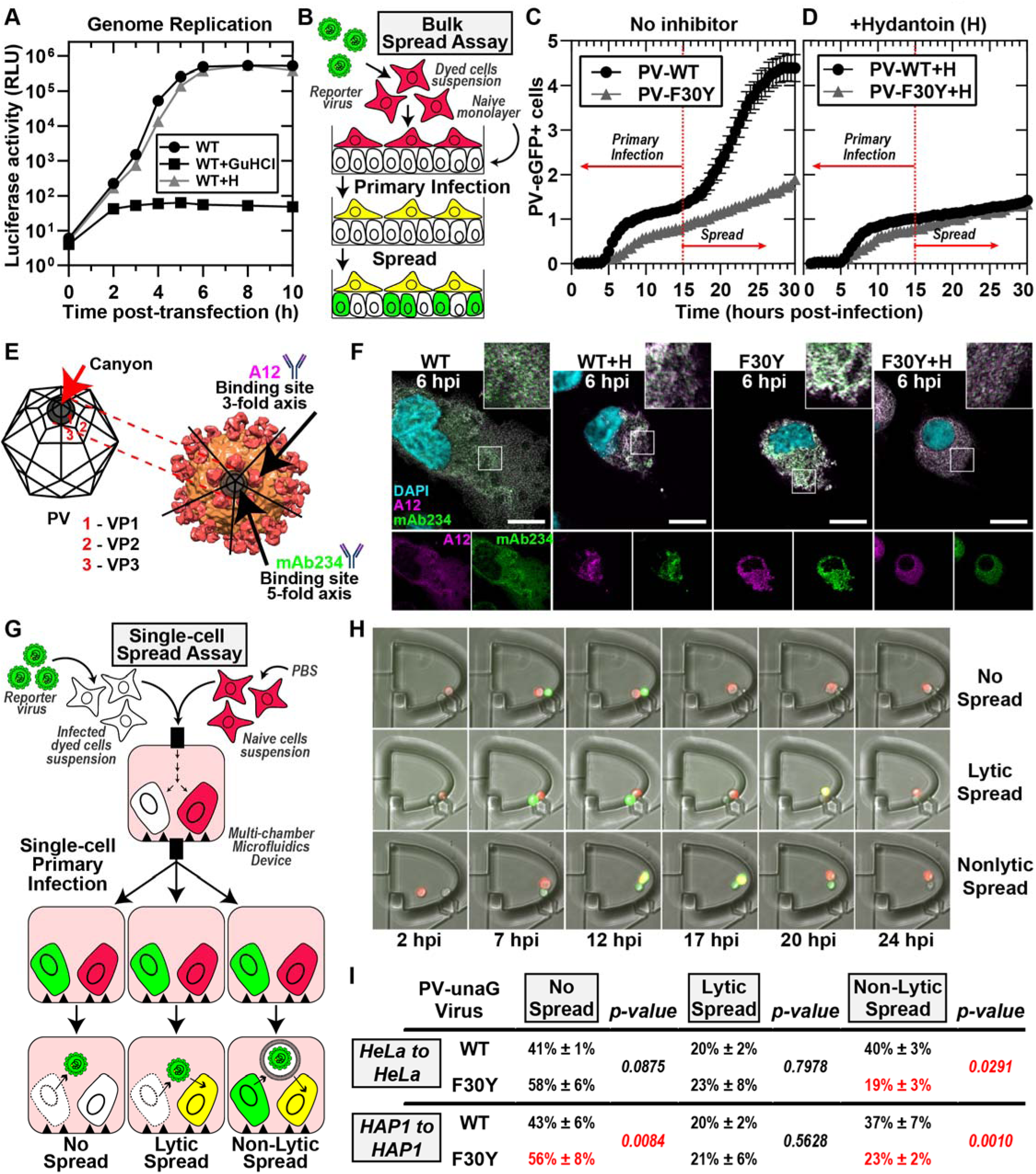
PV 3CD contributes to virion morphogenesis and non-lytic spread. **(A) PV genome replication in the presence of GuHCl and hydantoin.** PV sub-genomic replicon luciferase assay. HeLa cells were transfected with a WT PV replicon. in the presence and absence of 3mM GuHCl or 50 µg/mL hydantoin. Luciferase activity was measured as a surrogate for genome replication by relative light unit (RLU) from the collected lysates at the stated times/conditions. **(B) Bulk spread assay schematic.** HeLa cells in suspension were stained using a membrane dye and infected with a green fluorescence PVeGFP**_pv_** reporter variant. MOI of 5-infected dyed cells (red) were washed and seeded on top of a naïve HeLa cell monolayer. Fluorescence is monitored over time to detect both primary and secondary infections. Primary infected cells were observed and depicted in yellow when green (eGFP expression) and red signal (cell dye) colocalized. Spread was detected when a secondary wave of PV green fluorescence signal (green only) originating from the newly infected monolayer of unstained cells was observed. **(C) PV eGFP_pv_ and F30Y PV eGFP_pv_ bulk spread.** The graph depicts the number of eGFP-positive cells in a bulk spread assay performed as described in panel **(B)**. Using WT virus, the initial infection led to a spread event that increased the number of eGFP-positive cells (originating from secondary infections) observed after 15 hpi when the naive monolayer expressed eGFP signal (spread). F30Y PV eGFP_pv_ inhibited spread, as observed from a lack of a secondary wave eGFP signal. The data was normalized for the respective WT and F30Y eGFP infectivities. **(D) Bulk spread assay assessing the impact of hydantoin on PV spread.** The graph depicts the number of eGFP-positive cells in a bulk spread assay performed in the presence of 50 µg/mL hydantoin. **(E) PV structure and A12/mAb234 antibody illustrations.** WT PV icosahedron (left) and structure (right) illustrations indicate A12 and mAb234 antibodies-specific binding. A12 binds at the denoted 3-fold axis at the intersection of VP1, VP2, and VP3. MAb234 binds at the 5-fold axis where the canonical “canyon” is located. **(F) Confocal immunofluorescence imaging of A12 and MAb234 in PV-infected HeLa cells.** Images illustrate representative immunofluorescence image fields of WT- and F30Y-infected HeLa cells (MOI of 10) in the presence and absence of hydantoin. Cells were fixed and immunostained under the labeled conditions 6 hours post-infection (hpi). Fixed cells were immunostained using specific A12 (magenta) and mAb234 (green) antibodies. DAPI-stained nuclei are shown (cyan). The top panels show A12, mAb234, and DAPI fluorescence overlays. The bottom single panels show A12 and mAb234 fluorescence separately. **(G) Single-cell spread assay schematic.** Cells in suspension infected with a reporter PV-unaG_pv_ virus variant (green). Infected cells were paired with stained uninfected cells (red) in isolated chambers of a multi-chamber microfluidics polyvinylidene fluoride (PVDF) device. In this study, this device was modified to harbor cell pairs. Fluorescence is monitored over time to detect an initial wave of infected cells expressing green fluorescence., yielding a yellow fluorescence overlay (see yellow cells). Spread was detected when a secondary wave of green fluorescence signal was observed in red-dyed cells, producing a colocalized yellow signal. Spread events were further extrapolated into no-spread, lytic spread, and non-lytic spread. In no spread, no secondary infection signal was detected after a primary cell green fluorescence signal. In lytic spread, the secondary infection signal arose after losing the primary cell green fluorescence (lysis). In non-lytic spread, the secondary infection signal was detected while green fluorescence was still present in the primary infected cell. **(H) Epifluorescence imaging of single-cell pairs.** Representative fluorescence images of chambers harboring cell pairs in a single-cell spread assay. The panels describe each spread scenario described in **(G). (I) WT and F30Y unaG_pv_ single cell spread assay.** In this single-cell spread assay, HeLa or HAP1 cells were infected with either WT or F30YunaG_pv_ at an MOI of 5 and paired with uninfected stained cells (red). No spread, lytic, and non-lytic events are quantified as percentages of the total number of events. The values are represented as mean ± standard error (SEM) from an n=3. Significant differences between conditions were noted based on a student’s t-test with p-values below 0.05.

To assess PV spread at the population level, we use a recombinant GFP-expressing PV (49–51) to infect HeLa cells containing a plasma membrane red dye and place these on a monolayer of unstained, uninfected HeLa cells (Fig. 2B). As the primary infection proceeds, we observe the formation of yellow cells (**Fig. 2B**). As the virus spreads from the yellow cells to the uninfected cells, these cells appear green (**Fig. 2B**). We have provided a representative movie of PV spread using this assay (**Movie S1**).

We monitored the number of GFP-positive cells as a function of time for WT PV (PV-WT in **Fig. 2C**). We observed two phases. The first phase included a lag followed by linear accumulation of GFP-positive cells, with the rate of accumulation approaching a plateau by 15 hours post-infection (hpi) (PV-WT in **Fig. 2C**). At 15 hpi, a secondary wave of GFP-positive cells accumulated, creating an inflection point that is presumably a reflection of PV spread (PV-WT in **Fig. 2C**). That this second phase represented PV spread was supported by the sensitivity of this phase to the presence of hydantoin in the media (PV-WT+H in **Fig. 2D**).

We performed the same experiment using F30Y PV. We observed a linear accumulation of GFP-positive cells over the entire 30-h time course (PV-F30Y in **Fig. 2C**). The observed accumulation was not impacted by the presence of hydantoin (PV-F30Y+H in **Fig. 2D**), suggesting a slow rate of infection establishment and a substantial defect to and/or delay in virus assembly and/or virus spread.

To investigate virus assembly, we used two monoclonal antibodies: A12 (human) (52) and mAb234 (mouse) (53, 54). The epitope recognized by A12 is in the canyon and should be able to recognize assembled particles whether the viral genome has been encapsidated or not (**Fig. 2E**) (52, 55, 56). On the other hand, mAb234 binds to the rim of the canyon and should favor binding to a viral genome-containing particle, the virion (**Fig. 2E**) (54, 57, 58). We used these antibodies to assess infected cells at 6 hpi in the absence and presence of hydantoin (**Fig. 2F**). For WT PV, mAb234 exhibited the greatest reactivity, consistent with the presence of primarily infectious virions at this time point (WT in **Fig. 2F and Fig. S2A, C**). In the presence of hydantoin, however, A12 exhibited the greatest reactivity, consistent with hydantoin interfering with encapsidation of the viral genome to form infectious virions (WT+H in **Fig. 2F and Fig. S2A, C**).

Importantly, F30Y PV appears to have no problem making infectious virions based on the accumulation of mAb234-reactive virions as observed for WT (F30Y in **Fig 2F and Fig. S2B, C**). However, the localization of the virions appears to be restricted to the perinuclear regions compared to WT PV (F30Y in **Fig. 2F**). Hydantoin also interfered with the maturation of the virus produced by F30Y PV (F30Y+H in **Fig. 2F and Fig. S2B, C**).

We have developed a system to isolate cell pairs in nanowells and to monitor the spread of infection from an infected to an uninfected cell. This system will be described in detail in a separate report based on a previous design we published (51, 59). To distinguish primary from secondary infections, recipient uninfected cells were stained with a red dye. In these experiments, a UnaG reporter was used instead of GFP (**Fig. 2G**). The infected cell is introduced into a chamber with an uninfected cell, and infection dynamics are monitored in each cell by measuring green fluorescence evolution (**Fig. 2G**). A movie illustrating the experiment and outcomes has been provided (**Movie S2**). Three outcomes can be observed: no spread, lytic spread, and non-lytic spread (**Fig. 2G, H**). In a no-spread scenario, the infected cell lyses, but the released virus fails to establish infection in the uninfected cell.

We performed a single-cell-pairing experiment using two cell lines: HeLa and HAP1. Observations with each were essentially identical. The primary route of secondary infection for WT PV was by a non-lytic mechanism (non-lytic spread in **Fig. 2I**). The vast number of lytic infections for WT PV failed to result in secondary infections (compare no spread to lytic spread in **Fig. 2I**). Interestingly, F30Y PV was significantly and selectively impaired in its ability to spread by a non-lytic mechanism (**Fig. 2I**). Also interesting was the observation that loss of non-lytic infection led not to more lytic infections that spread but to more lytic infections that failed to spread (**Fig. 2I**).

### PV 3CD comigrates with virions from the perinuclear region of the cell to the periphery

Experiments shown in **Fig. 2F** above suggested a trafficking defect for virions produced by F30Y PV. We investigated this possibility directly, as indicated in **Fig. 3A**. We infected HeLa cells in the absence or presence of hydantoin and used immunofluorescence to monitor virus particles/virions and 3CD as a function of time post-infection. The specificity of the antibodies used is indicated by the absence of staining in mock-infected cells (**Fig. 3B**).

**Figure 3.**
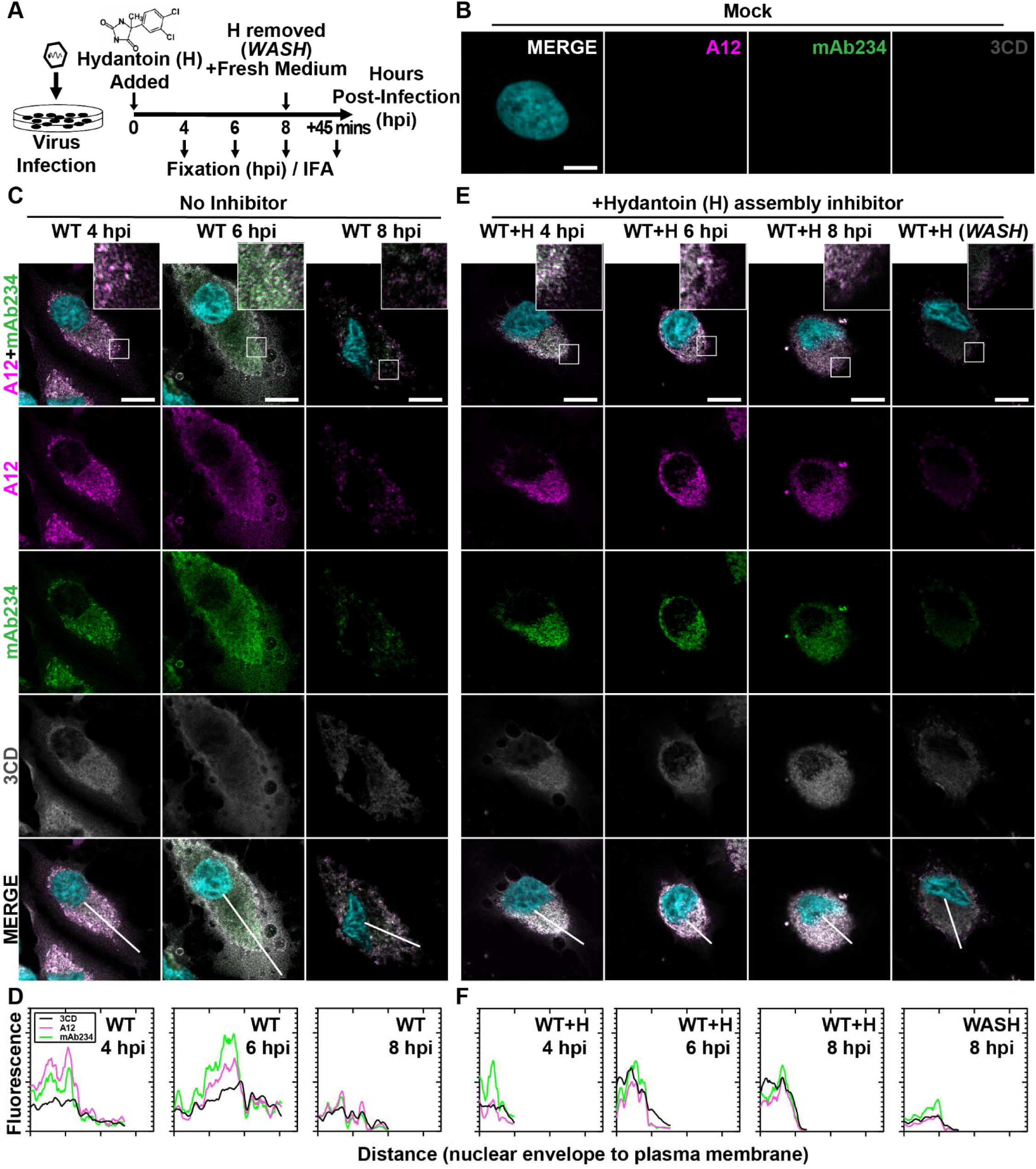
PV 3CD comigrates with virions from the perinuclear region of the cell to the periphery. **(A) PV time-course immunofluorescence assay (IFA) schematic.** HeLa cell monolayer infections were carried out in the presence or absence of hydantoin. Infected cells were fixed at the stated time points (4-, 6-, or 8-) hours post-infection. An additional timepoint labeled “WASH” was collected for cells undergoing 8 hours of infection in the presence of hydantoin; the monolayer was then rinsed with PBS to remove the drug. After rinsing, fresh, warm, complete medium was added, and cells were incubated for 45 minutes before fixing. An immunostaining fluorescence assay (IFA) was then conducted on fixed cells. **(B) Mock cell IFA.** Representative confocal immunofluorescence images of mock HeLa cells showing no virus A12, mAb234, or 3CD protein reactivity in the absence of PV infection. Uninfected cells were fixed and immunostained 6 hours after initiating the experiment. Fixed cells were immunostained using A12 (magenta), mAb234 (green), and 3CD (grey) antibodies. The DAPI-stained nucleus is shown (cyan). Overlays of all four fluorescence signals (MERGE) are shown. **(C) WT PV time-course IFA.** Images illustrate representative confocal immunofluorescence fields of WT-infected HeLa cells 4-, 6- and 8-hours post-infection (hpi). HeLa cells were infected with WT PV at an MOI of 10, fixed, and immunostained at the labeled time points. Fixed cells were immunostained as described for mock cells in panel **(B)**. The top panels show A12, mAb234, and DAPI fluorescence overlays with a perinuclear inset delineated with a white square. The bottom panels show A12, mAb234, 3CD, and DAPI fluorescence overlays (MERGE) with a white line extending from the nuclear envelope to the plasma membrane. Each column incrementally shows the hours post-infection from left to right 4-, 6-, and 8-hpi. **(D) WT PV fluorescence intensity profiles.** Intensity profile plots reveal the progression of A12, mAb234, and 3CD fluorescence over a WT PV infection time course. The bottom MERGE panels in **(C)** show a white line extending from the nuclear envelope to the plasma membrane used for “profile fluorescence” signal quantification. Intensity profile measurements were taken from regularly spaced points along a line segment to depict the spatial and temporal dynamics of fluorescence reactivity, levels, and signal overlap in infected cells over time. Values were plotted as a smooth line graph with relative fluorescence intensity units (RFU) on the Y-axis and distance (nm) on the X-axis. A12 (magenta), mAb234 (green), and 3CD (black) were plotted as independent lines in the graph. **(E) WT PV time-course IFA in the presence of hydantoin.** Images illustrate representative PV WT-infected HeLa cell confocal immunofluorescence fields in the presence of 50 µg/mL hydantoin (WT+H) 4-, 6-, and 8-hours post-infection (hpi) as described for WT in **(C).** An additional “WASH” time point indicates an infection where the hydantoin block is released at 8 hpi. **(F) WT PV fluorescence intensity profiles in the presence of hydantoin.** Intensity profile plots reveal the progression of A12, mAb234, and 3CD fluorescence over a WT PV infection time course in the hydantoin-inhibited state as described for WT in **(D)**. Intensity measurements were acquired from the WT+H panels shown in **(E)**.

At 4 hpi for WT PV, immature particles predominated (A12 staining exceeds mAb234 in WT 4 hpi column of **Fig. 3C**). 3CD protein was detected easily and colocalized with virus particles (MERGE in WT 4 hpi column of **Fig. 3C**). We quantified the staining by each antibody as a function of distance from the perinuclear region of the cell to the periphery (**Fig. 3D**). This analysis confirmed colocalization of 3CD proteins and particles (WT 4 hpi in **Fig. 3D**).

At 6 hpi for WT PV, virions predominated (mAb234 staining exceeds A12 in WT 6 hpi column of **Fig. 3C**). 3CD continued to colocalize with virions (MERGE in WT 6 hpi column of **Fig. 3C**). However, the peak of fluorescence for both virions and 3CD shifted away from the perinuclear region of the cell towards the periphery (WT 6 hpi in **Fig. 3D**). By 8 hpi for WT PV, most of the virions were no longer in the cell (WT 8 hpi column in **Fig. 3C**). Any residual staining of virus particles/virions occurred in the perinuclear regions of the cell (WT 8 hpi in **Fig. 3D**). The level of 3CD observed in the cell at 8 hpi was also diminished (compare 3CD at 6 hpi to 8 hpi in **Fig. 3C**).

### Immature virus particles do not move to the cell periphery

Hydantoin treatment is thought to trap an intermediate during genome encapsidation (46, 47), likely in association with the replication organelle (21). In the presence of hydantoin, immature particles accumulate and remain associated with the perinuclear region of the infected cell over the entire eight-hour period observed (A12 and mAb234 in columns WT+H 4, 6, 8 hpi of **Fig. 3E**). 3CD protein colocalized with virus particles and also failed to move to the periphery of the cell over the entire eight-hour period observed (3CD and MERGE in columns WT+H 4, 6, 8 hpi of **Fig. 3E**). Interestingly, removal of hydantoin followed by fixation 45 min later demonstrated a synchronous exodus of virions and some 3CD as well (WASH in **Fig. 3E**; compare WT+H 8 hpi to WASH in **Fig. 3E**).

### PV 3CD contributes to virion trafficking

We evaluated F30Y PV (**Fig. 4**) using the same series of experiments described immediately above for WT PV (**Fig. 3**). Despite the accumulation of mature virions, there is no detectable movement of the virions or 3CD from the perinuclear region to the periphery of the cell (**Figs. 4A, B**). The trafficking defect observed is similar to that observed in the presence of hydantoin (compare **Fig. 4A** to **Fig. 4C**). Unlike observed for WT PV, hydantoin block release did nothing to synchronize events related to egress (WASH in **Fig. 4C**). The failure of 3CD to leave the cell in the absence (3CD F30Y 8 hpi of **Figs. 4A, B**) or in the presence (3CD F30Y+H 8 hpi and WASH of **Figs. 4C, D**) of hydantoin suggest that it is not the mere duration of infection and corresponding cytopathic effect permeabilizing the cell to permit release of virions and/or 3CD protein.

**Figure 4.**
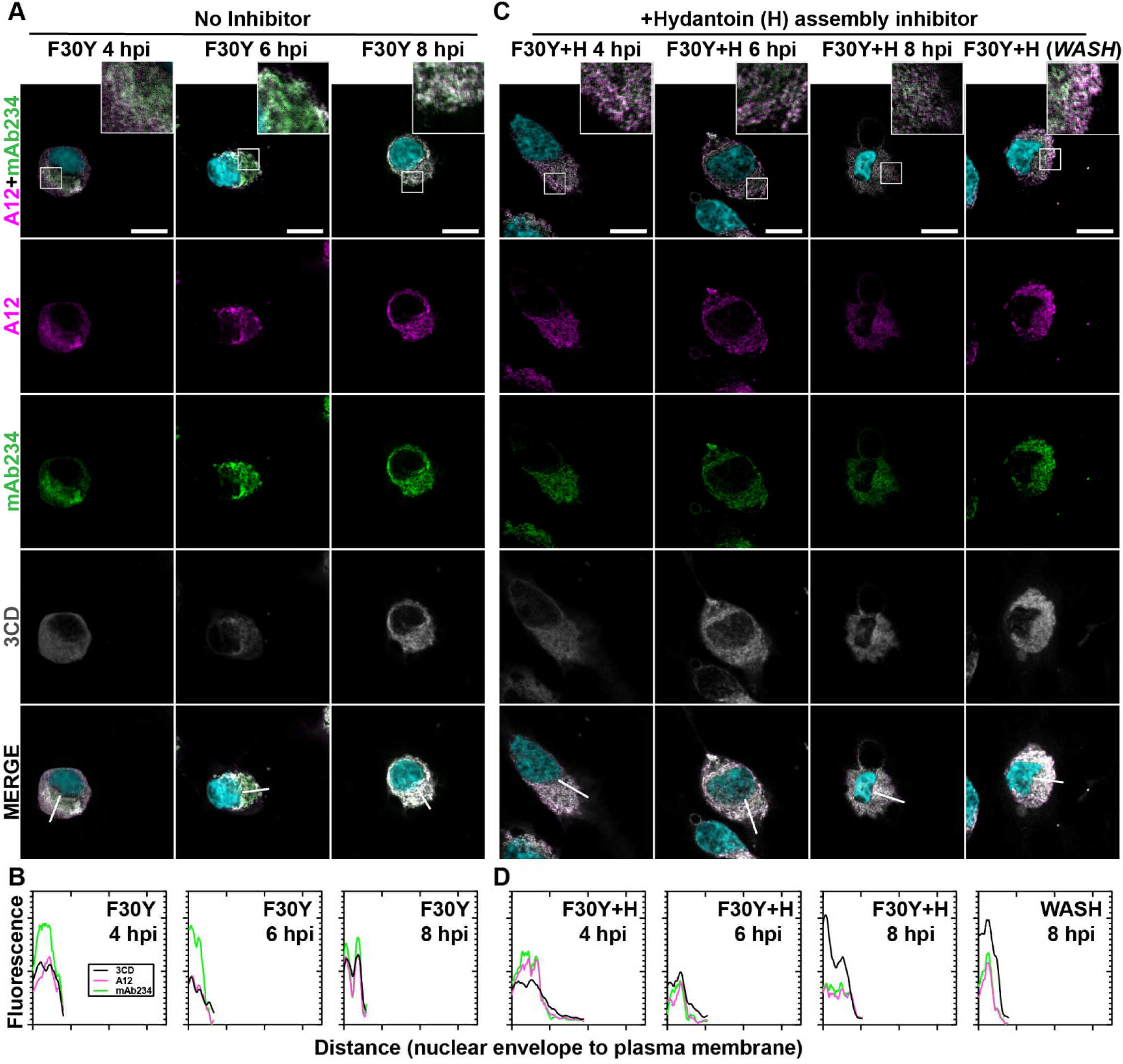
PV 3CD contributes to virion trafficking. **(A) F30Y PV time-course IFA.** Images illustrate representative confocal immunofluorescence fields of F30Y PV-infected HeLa cells 4-, 6- and 8-hours post-infection (hpi). HeLa cells were infected with F30Y PV at an MOI of 10, fixed, and immunostained at the labeled time points. Fixed cells were immunostained using A12 (magenta), mAb234 (green), and 3CD (grey) antibodies. DAPI-stained nuclei are shown (cyan). The top panels show A12, mAb234, and DAPI fluorescence overlays with a perinuclear inset delineated with a white square. The bottom panels show A12, mAb234, 3CD, and DAPI fluorescence overlays (MERGE) with a white line extending from the nuclear envelope to the plasma membrane. Each column shows hours post-infection incrementally from left to right 4-, 6-, and 8-hpi. **(B) F30Y PV fluorescence intensity.** Intensity profile plots reveal the progression of A12, mAb234, and 3CD fluorescence over an F30Y PV infection time course. The bottom panels in **(A)** show A12, mAb234, 3CD, and DAPI fluorescence overlays (MERGE), with a white line extending from the nuclear envelope to the plasma membrane used for “profile fluorescence signal” quantification. Intensity profile measurements were taken from regularly spaced points along a line segment to depict the spatial and temporal dynamics of fluorescence reactivity, levels, and signal overlap in infected cells over time. Values were plotted as a smooth line graph with relative fluorescence intensity units (RFU) on the Y-axis and distance (nm) on the X-axis. A12 (magenta), mAb234 (green), and 3CD (black) were displayed as independent values in the graph. **(C) F30Y PV time-course IFA in the presence of hydantoin.** Images illustrate representative confocal immunofluorescence fields of F30Y PV-infected HeLa cells in the presence of 50 µg/mL hydantoin (F30Y+H) as described in **(A)** for F30Y PV. An additional “WASH” time point indicates an infection where the hydantoin block is released at 8 hpi. **(D) F30Y PV fluorescence intensity in the presence of hydantoin.** Intensity profile plots reveal the progression of A12, mAb234, and 3CD fluorescence over an F30Y PV infection time course in the hydantoin-inhibited state. as described in **(B)** for F30Y PV. Intensity measurements were acquired from the F30Y+H panels shown in **(C)**.

### PV 3CD is required for colocalization of PV virions with lipidated LC3B

It is becoming increasingly clear that non-enveloped viruses like PV, other enteroviruses, and even more distantly related picornavirus family members move from one cell to another by non-lytic mechanisms (14, 16, 60). For PV and the other enteroviruses, secretory autophagy represents the most likely pathway used for virus spread (19–21, 61). For cellular homeostasis, this mechanism requires the correct cellular circumstance to exist for activation of a signaling cascade that leads to the formation of the site of assembly of autophagic vesicles, the so-called omegasome (**Fig. 5A**) (62–64). Cargo is recruited to the omegasome by a combination of factors, including microtubule-associated protein 1B-light chain 3 (LC3B) and adaptor proteins like sequestosome (SQSTM1/p62), which promotes even more selective cargo loading (65–68). Modification of LC3B with phosphatidylethanolamine produces a lipidated form referred to as LC3B-II (**Fig. 5B**). LC3B-II located within the omegasome recruits cargo proteins containing LC3-interacting regions (LIRs) (**Fig. 5C**). Cargo loading elongates the omegasome into a larger double-membraned structure that is ultimately sealed to produce an autophagosome (**Fig. 5A**) (62, 63, 66, 69–72). In secretory autophagy, autophagosomes move to the periphery of the cell, where they can fuse with the plasma membrane to release cargo enclosed within a single-membrane vesicle (**Fig. 5D**)(73–75).

**Figure 5.**
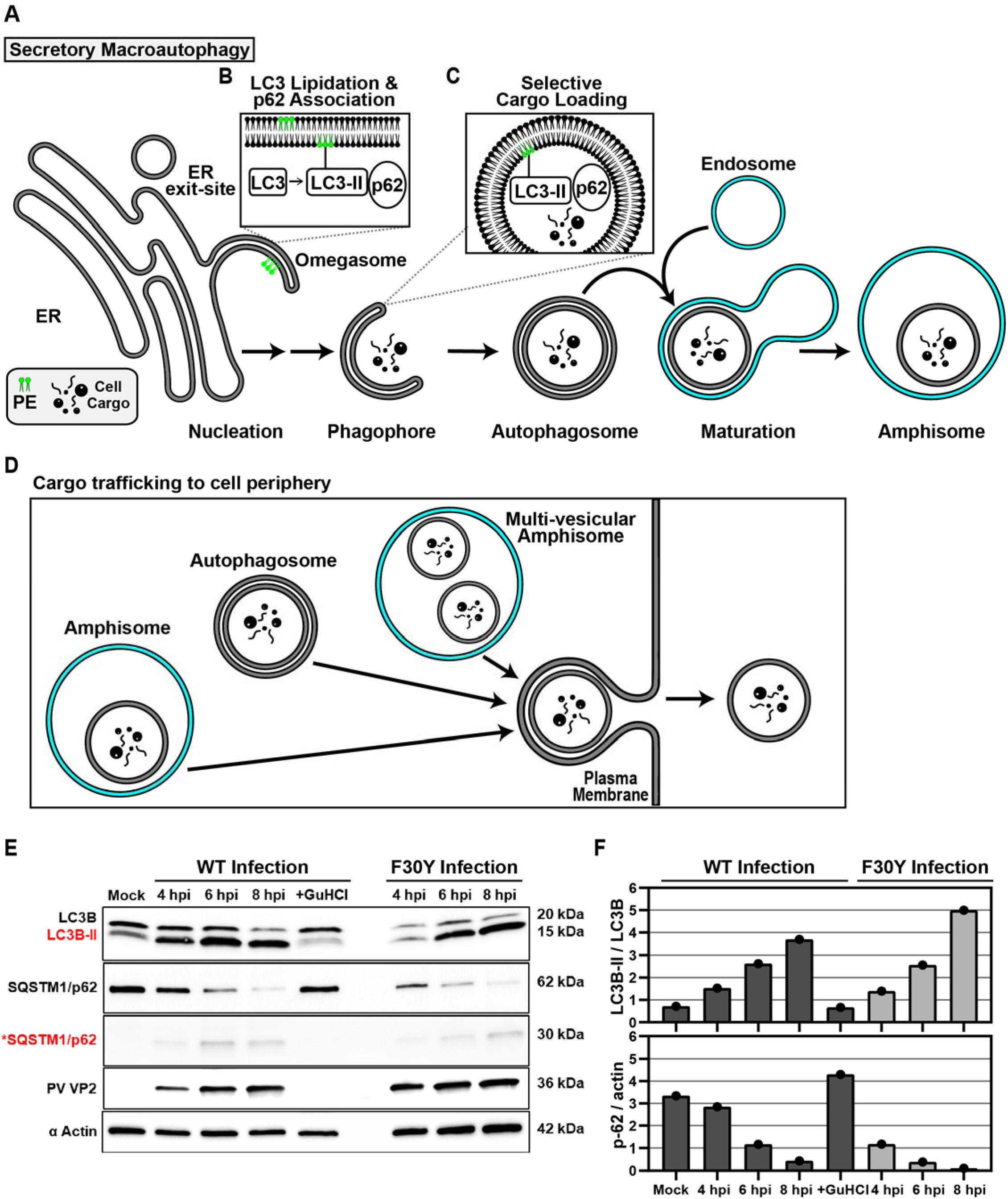
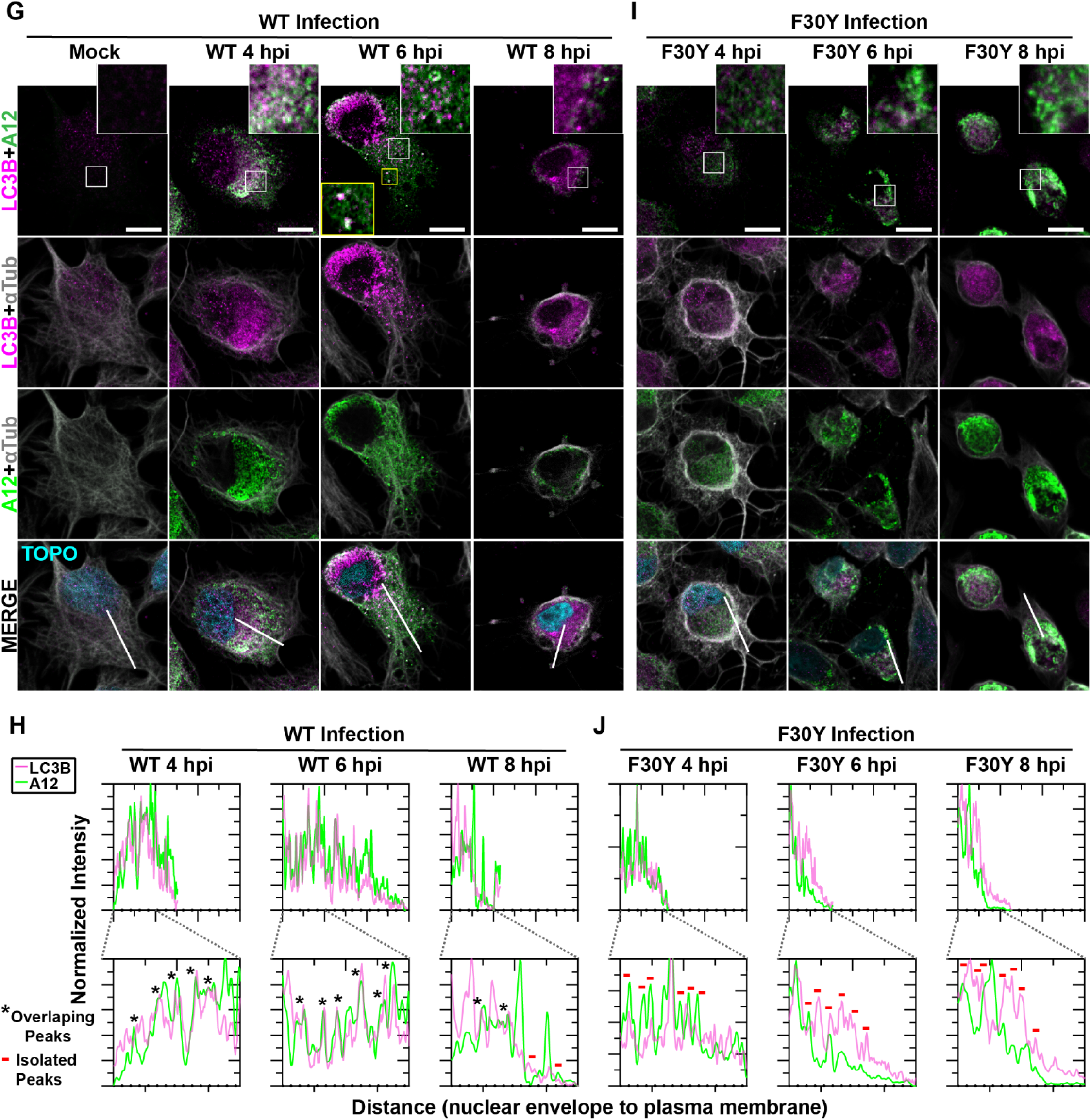
PV 3CD is required for colocalization of PV virions with lipidated LC3B. **(A) Autophagy pathway schematic.** An ER-derived omegasome buds out and is engaged by multiple autophagy-associated proteins, adaptors, kinases, and protein complexes to yield an autophagophore in preparation for cargo loading and maturation of a double-membranous vesicle termed autophagosome. **(B)** Autophagosome maturation is triggered by the lipidated form of the essential microtubule-associated protein 1A/1B-light chain 3 (LC3) protein. **(C)** Cargo is recruited to the phagophore by combining factors, including LC3 and adaptor proteins like sequestosome (SQSTM1/p62), which promote selective cargo loading. For intact/functional cargo secretion in vesicles, the autophagosome may fuse with endosomes to form a cargo-containing amphisome. **(D)** Cargo-containing amphisomes, multi-vesicular amphisomes, and/or autophagosomes can then be trafficked to the plasma membrane and secreted onto the extracellular space. **(E) WT and F30Y PV infection immunoblots.** Images show representative immunoblots of WT and F30Y PV-infected cell lysates. Cells were infected with the indicated conditions, and lysates were collected at the displayed time points 4-, 6-, and 8-hours post-infection (hpi). Both mock and GuHCl control for infection and genome replication phenotypes, respectively. Lysates were then subject to western blot analysis and probed with LC3B, SQSTM1/p62, PV VP2, and α actin antibodies. **(F) LC3B lipidation and SQSTM1/p62 cleavage quantification.** WT and F30Y PV infection immunoblot quantification of LC3B and SQSTM1/p62 chemiluminescence signals. The ratio of lipidated-LC3B protein (LC3B-II) to LC3B protein increases while the full-length SQSTM1/p62 protein levels decrease as the infection progresses in WT and F30Y PV-infected HeLa cells. **(G) WT PV time course – LC3B IFA.** Images illustrate representative confocal immunofluorescence fields of WT-infected HeLa cells 4-, 6- and 8 hours post-infection (hpi). HeLa cells were infected with WT PV at an MOI of 10, fixed, and immunostained at the labeled time points. Fixed cells were immunostained using LC3B (magenta), A12 (green), and αTubulin (grey) antibodies. TOPO-stained nuclei are shown (cyan). The top panels show LC3B and A12 fluorescence overlay with a perinuclear inset delineated with a white square and a cytoplasmic inset in yellow. The bottom panels show LC3B, A12, αTubulin, and TOPO fluorescence overlays (MERGE) with a white line extending from the nuclear envelope to the plasma membrane of cells. Each column incrementally shows the hours post-infection from left to right mock, 4-, 6-, and 8-hpi. **(H) WT PV fluorescence intensity profiles.** Intensity profile plots reveal the progression of LC3B and A12 fluorescence in WT PV-infected cells over time. The bottom panels in **(G)** show LC3B, A12, αTubulin, and TOPO fluorescence overlays (MERGE), with a white line extending from the nuclear envelope to the plasma membrane used for “profile” fluorescence signal quantification. Intensity profile measurements were taken from regularly spaced points along a line segment to depict the spatial and temporal dynamics of fluorescence reactivity, levels, and signal overlap in infected cells over time. Values were plotted as a smooth line graph with relative fluorescence intensity units (RFU) on the Y-axis and distance (nm) on the X-axis. LC3B (magenta) and A12 (green). **(I) F30Y PV time course – LC3B IFA.** Images illustrate representative confocal immunofluorescence fields of F30Y PV-infected HeLa cells 4-, 6-, and 8 hours post-infection (hpi). Fixed cells were immunostained using LC3B (magenta), A12 (green), and αTubulin (grey) antibodies as described for WT PV in **(G)**. TOPO-stained nuclei are shown (cyan). **(J) F30Y PV fluorescence intensity profiles.** Intensity profile plots reveal the progression of LC3B and A12 fluorescence of F30Y PV-infected cells over time. Intensity measurements were acquired from the panels shown in **(I)** as described for WT PV in **(H)**.

We hypothesized that the ability of particles to be loaded into autophagosomes was impaired for F30Y PV. From qualitative (**Fig. 5E**) and quantitative (**Fig. 5F**) perspectives, none of the early steps of PV-induced autophagic signals were changed for the mutant virus, including LC3B lipidation or SQSTM1/p62 cleavage. The use of GuHCl permitted assessing virus-induced changes triggered by proteins produced during the pioneer round(s) of translation of the viral genome (+GuHCl in **Figs. 5E, F**) (44, 45).

By monitoring the colocalization of virions with LC3B-II as a function of time post-infection, we observed a strong colocalization for WT PV (**Fig. 5G**). There was a strong correspondence in fluorescence intensity patterns from the perinuclear region to the periphery of the cell for virions and LC3B-II, consistent with virions being moved to the cell periphery in autophagosomes or more complex structures, for example, amphisomes (**Fig. 5H**) (62, 75–77). In contrast, the colocalization of virions with LC3B-II for F30Y PV was, at best, weak, if it occurred at all, either at the level of overlap of the intracellular fluorescence (Fig. 5I) or when monitoring the pattern of fluorescence intensity (**Fig. 5J**).

### Virions produced by both WT and F30Y PV colocalize with GABARAP

LC3 paralogues are collectively referred to as mammalian autophagy-related 8 (Atg8) family members, with Atg8 referring to the yeast orthologue. One paralogue implicated in non-lytic spread by PV is GABA Type A Receptor-Associated Protein (GABARAP), but insufficient data exist to attribute such a function to GABARAP definitively (21, 71, 78).

For WT PV, virions and GABARAP colocalized over the timeframe evaluated and moved from the perinuclear region to the periphery of the cell (**Fig. 6A**). The observed colocalization was confirmed by quantitative evaluation of the pattern of fluorescence intensity (**Fig. 6B**). While F30Y PV caused accumulation of both virions and GABARAP in the perinuclear region (**Fig. 6C**), colocalization of these proteins was not reduced substantially in the presence of the 3CD derivative (**Fig. 6D**).

**Figure 6.**
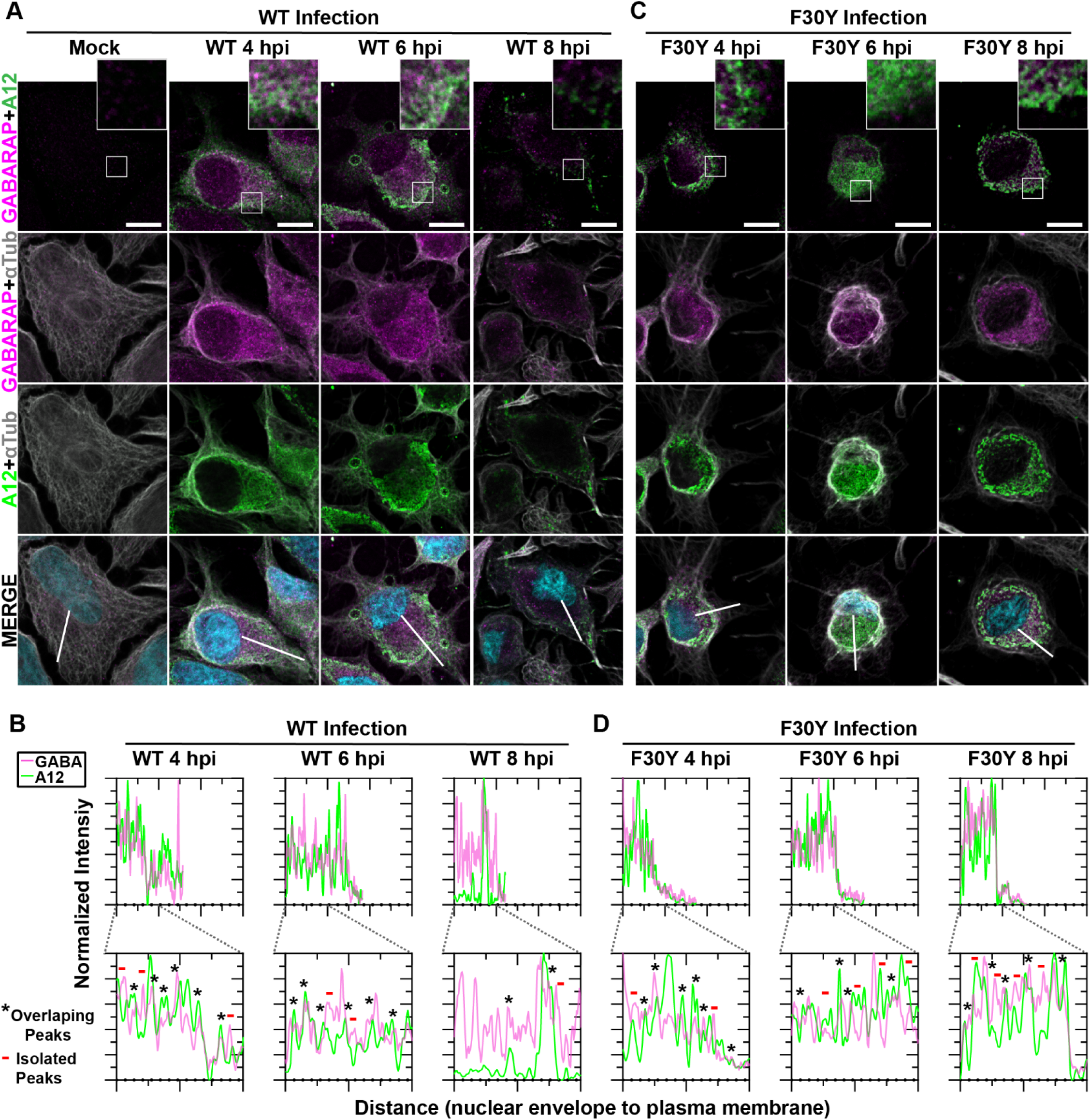
Virions produced by both WT and F30Y PV colocalize with GABARAP. **(A) WT PV time course – GABARAP IFA.** Images illustrate representative confocal immunofluorescence fields of WT-infected HeLa cells 4-, 6- and 8 hours post-infection (hpi). HeLa cells were infected with WT PV at an MOI of 10, fixed, and immunostained at the labeled time points. Fixed cells were immunostained using GABARAP (magenta), A12 (green), and αTubulin (grey) antibodies. TOPO-stained nuclei are shown (cyan). The top panels show GABARAP and A12 fluorescence overlay with a perinuclear inset delineated with a white square and a cytoplasmic inset in yellow. The bottom panels show GABARAP, A12, αTubulin, and TOPO fluorescence overlays (MERGE) with a white line extending from the nuclear envelope to the plasma membrane of cells. Each column incrementally shows the hours post-infection from left to right mock, 4-, 6-, and 8-hpi. **(B) WT PV fluorescence intensity profiles.** Intensity profile plots reveal the progression of GABARAP and A12 fluorescence in WT PV-infected cells over time. The bottom panels in **(A)** show GABARAP, A12, αTubulin, and TOPO fluorescence overlays (MERGE), with a white line extending from the nuclear envelope to the plasma membrane used for “profile” fluorescence signal quantification. Intensity profile measurements were taken from regularly spaced points along a line segment to depict the spatial and temporal dynamics of fluorescence reactivity, levels, and signal overlap in infected cells over time. Values were plotted as a smooth line graph with relative fluorescence intensity units (RFU) on the Y-axis and distance (nm) on the X-axis. GABARAP (magenta) and A12 (green). **(C) F30Y PV time course – GABARAP IFA.** Images illustrate representative confocal immunofluorescence fields of F30Y PV-infected HeLa cells 4-, 6-, and 8 hours post-infection (hpi). Fixed cells were immunostained using GABARAP (magenta), A12 (green), and αTubulin (grey) antibodies as described for WT PV in **(A)**. TOPO-stained nuclei are shown (cyan). **(D) F30Y PV fluorescence intensity profiles.** Intensity profile plots reveal the GABARAP and A12 fluorescence progression of F30Y PV-infected cells over time. Intensity measurements were acquired from the panels shown in **(I)** as described for WT PV in **(C)**.

### Application of high-angle annular dark-field (HAADF) scanning transmission electron microscopy (STEM) to the study of PV-induced autophagic signals

As discussed above, PV non-lytic spread almost certainly hijacks the secretory autophagy pathway (19). However, conventional transmission electron microscopy (TEM) has yet to yield high-resolution images of membrane-associated or vesicular intermediates, no matter how the cells are processed (8, 22, 79). We have used high-angle annular dark-field (HAADF) scanning transmission electron microscopy (STEM) on a 200 kV instrument. HAADF STEM affords good mass contrast without standard TEM staining (**Fig. S1**).

We focused first on analyzing PV-infected cells at 6 hpi, a time in which virion morphogenesis should be complete and trafficking of particles to the cell periphery should be in progress. In contrast to conventional TEM, electron density appears white in images collected using HAADF STEM. Even at the lowest magnification, we observed vesicular structures filled with electron density (1 μm in **Fig. 7A**) that improved in resolution as the magnification increased (500 nm and 100 nm in **Fig. 7A**). At the highest magnification, we observed double-membrane vesicles containing virus particles. By surveying multiple cells, we collected images consistent with steps of the secretory autophagy pathway (**Fig. 7B**) as illustrated in **Fig. 7C**: omegasomes, autophagosomes, and amphisome-like vesicles containing virions.

**Figure 7.**
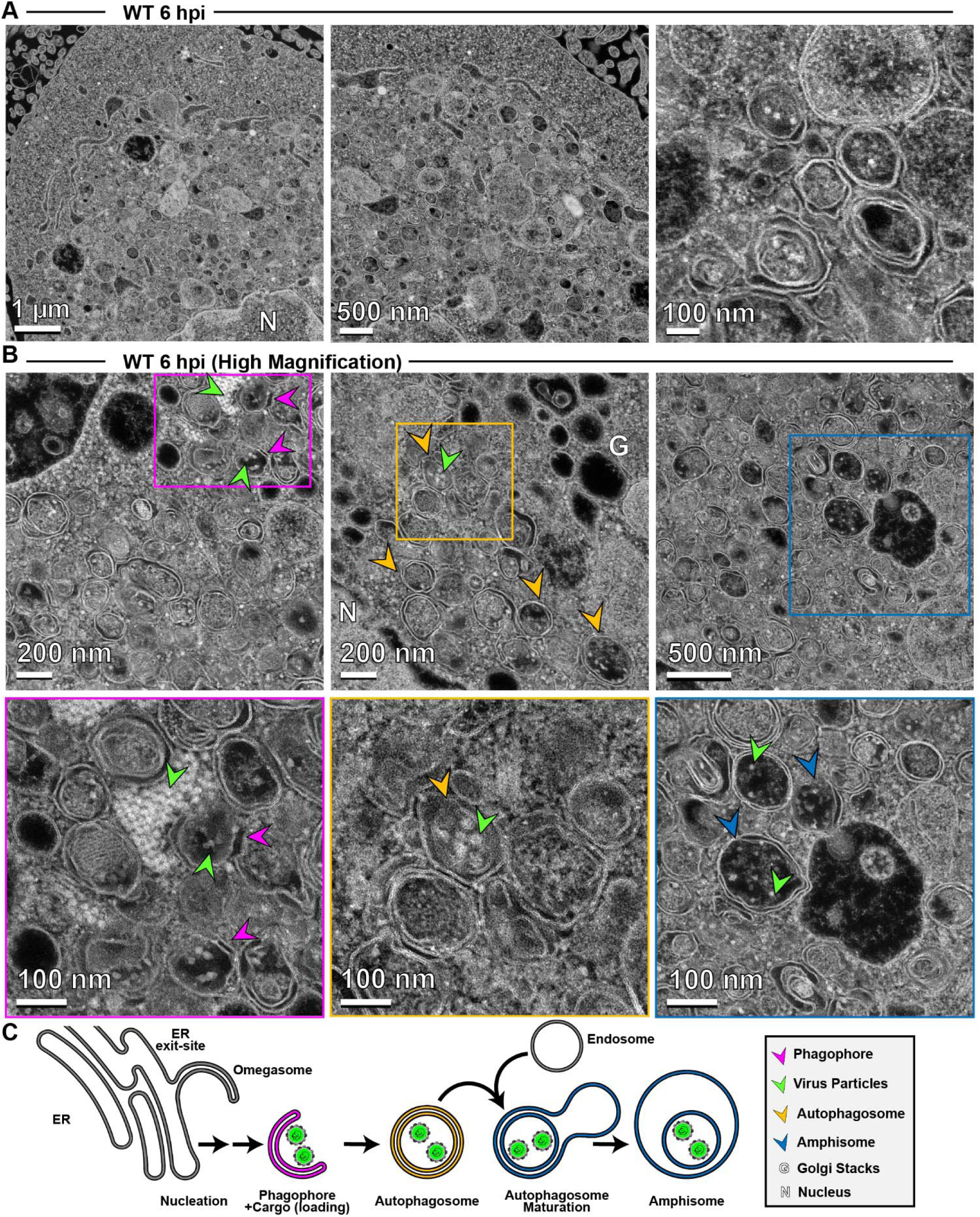
Application of high-angle annular dark-field (HAADF) scanning transmission electron microscopy (STEM) to the study of PV-induced autophagic signals. **(A) HAADF-STEM imaging of WT PV-infected HeLa cells.** HeLa cells were infected with WT PV at an MOI of 10 and then fixed in glutaraldehyde at the indicated time points. Fixed samples were dehydrated, stained, embedded, and sectioned in thin micrographs for imaging as described (**Fig. S1**). Images were collected using a Thermo Scientific Talos F200X G2 (S)TEM operated at 200 kV and a beam current of approximately 0.12 nA. The contrast is also reversed when compared to TEM, with the vacuum appearing dark. WT infection induces virus-containing double membranous vesicles and amphisome-like vesicles with virions in the intra-luminal vesicles. Arrows indicate observed structures. Phagophore (magenta), virus particles (green), autophagosomes (yellow), amphisomes (blue), Golgi (G), nucleus (N). Large outer vesicles with intra-luminal vesicles (100-300 nm diameter) contain ∼30 nm particles inside. Double membrane vesicles are located at sites where vesicular-tubular clusters are observed in TEM mode (see **Fig. S1**). **(B) STEM imaging of WT PV-infected HeLa cells** (**Magnified).** In this magnified view of STEM images, 30 nm virus particles were observed inside intra-luminal vesicles. Close-up view of an intra-luminal vesicle that contains 30 nm particles. **(C) Autophagic signals during WT PV infection.** An ER-derived omegasome buds out and is engaged by multiple autophagy-associated proteins, adaptors, kinases, and protein complexes to yield an autophagophore in preparation for virion loading and maturation of a double-membranous vesicle termed autophagosome. For intact/functional cargo secretion in vesicles, the autophagosome may fuse with endosomes to form a virus-containing amphisome-like vesicle.

In the presence of hydantoin, omegasomes and autophagosomes formed (**Fig. 8A**). However, these structures were either devoid of cargo based on the absence of electron density or contained an unknown fibrous material that did not exhibit strong electron density (**Fig. 8A**). These latter structures were also visible during normal infection in the absence of hydantoin (**Fig. 7B**).

**Figure 8.**
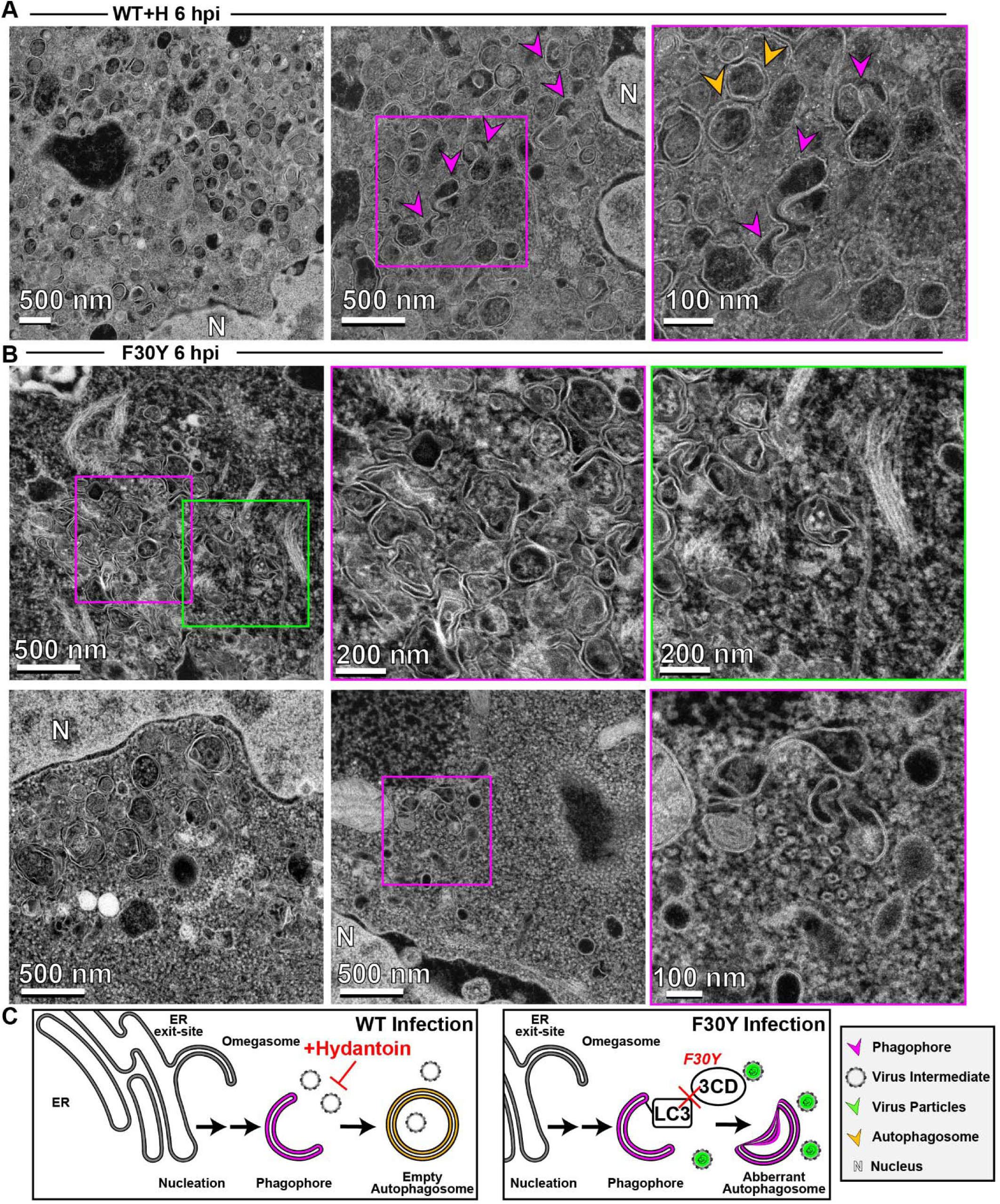
3CD-mutant PV exhibits defects to autophagosome biogenesis and virion loading. **(A) HAADF-STEM imaging of WT PV-infected HeLa cells in the presence of hydantoin.** HeLa cells were infected with WT PV at an MOI of 10 in the presence of 50 µg/mL hydantoin and then fixed in glutaraldehyde at the indicated time points. Fixed samples were dehydrated, stained, embedded, and sectioned in thin micrographs for imaging as described (**Fig. S1**). Arrows indicate observed structures. Phagophore (magenta), virus particles (green), autophagosomes (yellow), and nucleus (N). Hydantoin impairs virus assembly, as evidenced by the lack of virus particles observed in the image. Omegasomes, empty double-membrane vesicles (DMVs), and fiber-like structure-containing DMVs are abundant in these samples. Intra-luminal vesicles in amphisome-like vesicles appear empty. These ultrastructural changes are observed both at 6 and 8 hpi. **(B) HAADF-STEM imaging of F30Y PV-infected HeLa cells.** HeLa cells were infected with F30Y PV at an MOI of 10 and then fixed in glutaraldehyde at the indicated time points. Fixed samples were dehydrated, stained, embedded, and sectioned in thin micrographs for imaging as described (**Fig. S1**). Arrows indicate observed structures. Phagophore (magenta), virus particles (green), autophagosomes (yellow), and nucleus (N). F30Y interferes with DMV maturations with an exaggerated amount of omegasomes and aberrant DMVs observed by 6 hpi. Few virions are observed, some of which appear “stuck” in an omegasome. **(C) Autophagic signals during F30Y PV infection.** An ER-derived omegasome buds out and is engaged by multiple autophagy-associated proteins, adaptors, kinases, and protein complexes to yield an autophagophore in preparation for virion loading and maturation of a double-membranous vesicle termed autophagosome. This step is blocked by hydantoin. Autophagosome maturation is triggered by the lipidated form of the essential microtubule-associated protein 1A/1B-light chain 3 (LC3) protein. Virus is recruited to the phagophore by combining factors, including LC3 and 3CD. F30Y 3CD interferes with this step.

Interestingly, F30Y PV exhibited a unique phenotype. Omegasomes accumulated (**Fig. 8B**). Few vesicular structures containing virions existed (**Fig. 8B**). When virions were loaded into omegasome-like structures, potentially derived from a replication organelle precursor, these failed to close and form autophagosomes (**Fig. 8B**). A large number of empty, single-membrane vesicles were also present in the perinuclear region of the cell (**Fig. 8B**).

Together, these observations suggest at least two independent functions of 3CD in autophagic vesicle formation used for non-lytic spread. First, 3CD contributes to omegasomes closure to form autophagosomes (**Fig. 8C**). Second, 3CD contributes to loading of virions into omegasomes/autophagosomes, likely in an LC3II-dependent manner (**Fig. 8C**). Both of these 3CD activities are impaired in the F30Y derivative. Further validating this claim is the observation that 3CD protein can be recovered from inside virus-containing vesicles isolated from the supernatant of virus-infected cells (80).

### LC3- and GABARAP-interacting regions in PV 3CD

The apparent loss of interaction between LC3B-II and virions in the presence of F30Y 3CD (**Fig. 5E**) prompted us to evaluate the presence of an LC3-interacting region (LIR) in 3CD. We used the iLIR Database algorithm to search for the LIR consensus: (W/F/Y)-(X)-(X)-(L/I/V) (81). We identified 13 putative LIRs in the 3C and 3D domains of 3CD (**Fig. 9A**). Interestingly, F30 defines the first amino acid of an LIR consensus motif, and this motif is conserved across enteroviruses (**Fig. 9B**).

**Figure 9.**
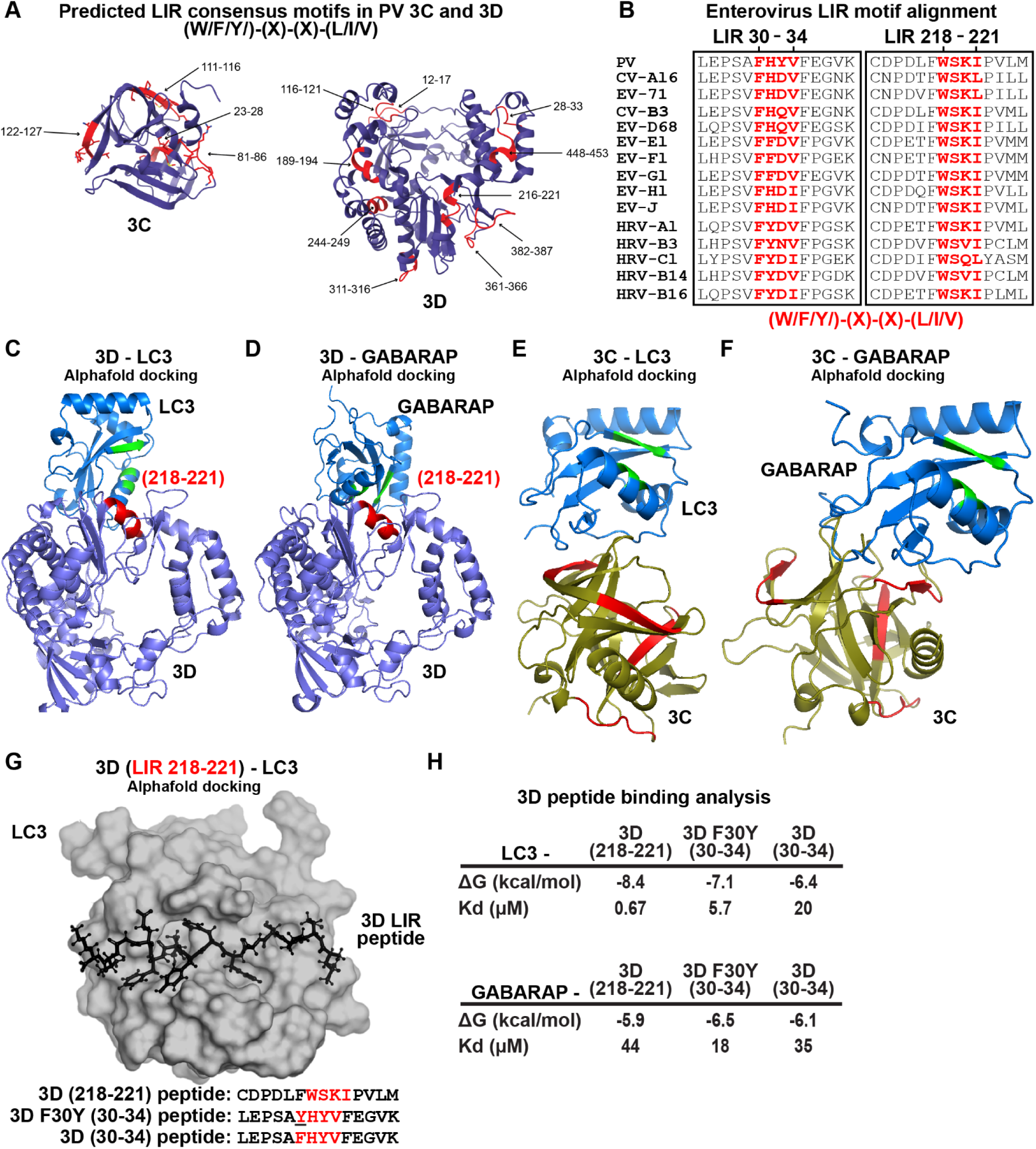
LC3- and GABARAP-interacting regions in PV 3CD. **(A) PV 3C and 3D LIRs.** LC3-interacting region (LIR) mediates LC3 binding with autophagy-associated factors and cargo. LIRs are characterized by a consensus motif (W/F/Y) (x) (x) (L/I/V). All PV protein products encode at least 1 LIR for a total of 33 across all PV proteins. The 3CD region encodes for 13 LIRs. Shown in violet are ribbon depictions of 3C (PDB 1L1N) and 3D (PDB 1RA6), with LIRs highlighted in red. **(B) Enterovirus 3D LIRmotif alignment.** Enteroviruses encode at least two W/F/Y) (x) (x) (L/I/V) LIRs in the 3D region, which are strictly conserved across multiple virus species. The panel represents a partial sequence alignment of the PV 3D “palm” and “thumb” subdomains. Two motif regions that follow the LIR consensus sequence pattern are highlighted in red. **(C) PV 3D and LC3A docking.** Alphafold docking of 3D (violet ribbon depiction with an LIR in red) with LC3A (blue ribbon depiction with hydrophobic pocket in green) **(D) PV 3D and GABARAP docking.** The panels represent Alphafold docking of 3D (violet cartoon with LIR in red) with GABARAP (blue cartoon with hydrophobic pocket in green). **(E) PV 3C and LC3B docking.** The panels represent Alphafold docking of 3C (olive cartoon with LIR in red) with LC3B (blue ribbon with hydrophobic pocket in green). **(F) PV 3C and GABARAP docking.** The panels represent Alphafold docking of 3C with GABARAP (blue cartoon with hydrophobic pocket in green). **(G) PV 3D LIR peptide and LC3B docking.** The panel represents Alphafold docking of the LEPSAF(30)HYVFEGVK peptide (in black ball-and-stick) with LC3B (gray surface). **(H) PV 3D and LC3B peptide binding analysis.** Analysis of docking and binding of two WT and mutant peptides to LC3B.

To ask whether or not any of the LIRs could interact with LC3 or GABARAP, we performed a computational docking experiment (82, 83). This experiment identified the region between 216-221 of 3D as capable of interacting with LC3 (**Fig. 9C**) and GABARAP (**Fig. 9D**) in the natively folded protein but not the region between 28-33.

Performing the same experiment with 3C revealed no interaction with LC3 (**Fig. 9E**) or GABARAP (**Fig. 9F**) mediated by a predicted LIR. However, an interaction was observed in these experiments. Assessment of the relevance of this interaction, if at all, will require additional experiments.

The preceding analysis failed to explain a defect associated with the F30Y substitution. We know that 3CD exhibits substantial conformational dynamics (26, 28, 84). We reasoned that a conformation may exist to permit an interaction between the 28-33 LIR of 3D that the F30Y substitution might disrupt. To test this possibility, we docked the 28-33 (WT and F30Y) and 216-221 peptides of 3D to the LIR-binding sites of LC3 and GABARAP (**Fig. 9G**) (82, 83). We calculated the corresponding values for their equilibrium dissociation constants (*K*_d_) (**Fig. 9H**). This analysis showed that 216-221 LIR bound more tightly to LC3 and GABARAP than the 28-33 LIR. However, the F30Y substitution increased affinity for LC3/GABARAP rather than decreased it.

As a final approach to obtain insight into the biochemical/biophysical basis for the defect associated with the F30Y substitution, we performed a molecular dynamics simulation of PV 3CD and the F30Y derivative (**Fig. 10**). PV 3CD is comprised of two domains separated by an interdomain linker and exhibits substantial conformational heterogeneity when comparing the orientation of one domain to that of the other (26, 28, 84). The 3C domain of the derivative was in a different location for the F30Y derivative (**Fig. 10A**). This difference is likely not meaningful. However, some of the more subtle conformational differences, reported as RMSD, may have some relevance (**Fig. 10B**).

**Figure 10.**
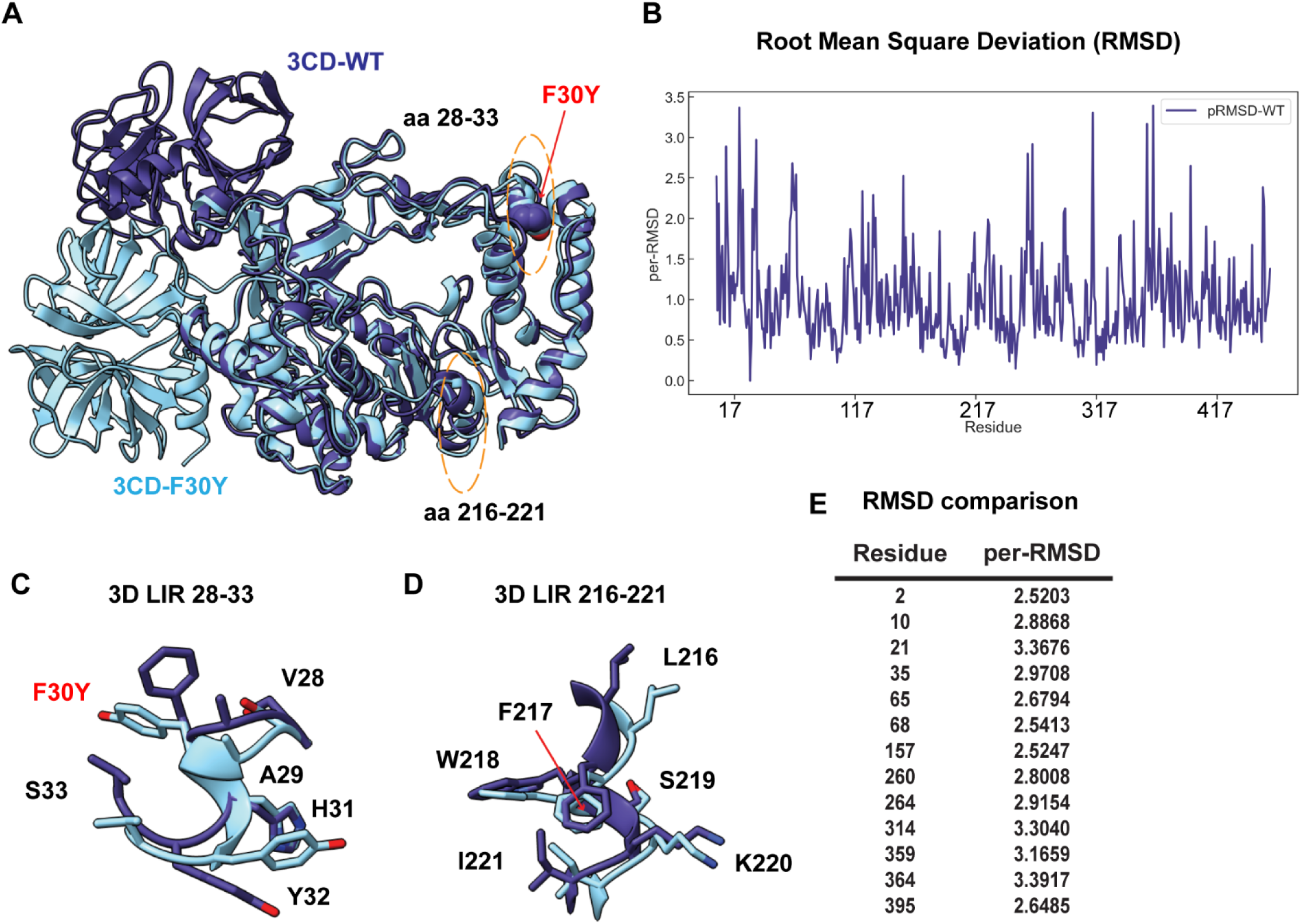
PV RdRp per-residue RMSD analysis suggests distinct conformational changes between WT and F30Y. **(A) MD-simulated WT and F30Y PV 3CD structures.** Depicted are the most visited conformations of WT (dark slate blue) and F30Y (cyan) structures from MD simulations that are superimposed and shown as cartoons. The predicted 3D (28-33) LIR locations with the F30Y variant highlighted in red and (217-221) LIR are highlighted in dotted yellow ovals. **(B) Root mean square deviation (RMSD)**. RMSD between the simulated WT and F30Y structures is plotted for the polymerase domain (aa 1-461, numbering corresponds to the 3D domain of 3CD). RMSD values were calculated using non-hydrogen atoms and averaged per-residue (per-RMSD). High per-residue RMSD values (>2.0) indicate regions of the polymerase that exhibited differences in conformations between WT and F30Y during MD simulations. **(C) 3D LIR** (**28-33**) **F30Y conformational changes.** Magnified view of the PV 3D (28-33) showing the distinct sidechain conformations of LIR residues 28-33. WT PV 3D is displayed in violet and F30Y in cyan. **(D) 3D LIR** (**217-221**) **F30Y conformational changes.** Magnified view of the PV 3D (217-221) showing the distinct sidechain conformations of LIR residues 217-221. WT PV 3D is displayed in violet F30Y in cyan. **(E) RMSD comparison.** Table describing highlighted values from the per-RMSD calculations between WT and F30Y PV 3CD with values higher than 2.0.

For example, the 28-33 LIR had a clear conformational difference (**Fig. 10C**) that did not occur with the 217-221 LIR (**Fig. 10D**). Moreover, residues across the entire protein exhibited significant changes in the 2.5 – 3.5 Å^2^ range because of the single Phe-to-Tyr change (**Fig. 10E**). Such changes suggest an allosteric connection between the 28-33 LIR and other regions of the protein that could be responsible for the phenotype observed here.

## DISCUSSION

It is now widely believed that dissemination of animal non-enveloped viruses from one cell to another or from one organ/tissue to another exploit virus-repurposed, vesicular carriers of the cell to produce "quasi-enveloped" or "vesicle-cloaked" virions (14, 16, 60, 85). Such a mechanism limits cell lysis and the enormous inflammatory response that would ensue if a lytic mechanism of spread were obligatory. This mechanism also increases the multiplicity of virions initiating an infection (86). Selective, secretory autophagy appears to be the primary mechanism used by enteroviruses (13, 19–21).

Since the earliest suggestion that autophagy contributes to enterovirus multiplication (87), many studies have focused on visualizing the double-membrane carriers in infected cells and evaluating the extent to which the *normal* cellular autophagy pathway and corresponding factors contribute to virus-induced autophagic signals (18, 21, 24, 88, 89). Only a few studies implicate an enteroviral protein(s) in the pathway enteroviruses use for non-lytic spread (24, 89, 90). Here, we report studies designed to probe the structure-function relationships governing processive RNA synthesis by the poliovirus (PV) RNA-dependent RNA polymerase (RdRp) that identified a derivative of the PV nonstructural protein 3CD that substantially reduced PV spread (**Fig. 1**), especially non-lytic spread (**Fig. 2**). The goal of this study therefore became focused on elucidating the step(s) post-genome replication requiring 3CD that ultimately leads to reduced, non-lytic spread.

The intracellular, post-genome-replication steps of the enterovirus lifecycle are the least understood. However, studies from several laboratories published over the past decade or so provide a framework for these late lifecycle steps when considered together (16, 19, 23). PV infection induces the formation of membrane tubules that are thought to support genome replication (8, 91, 92). As genome replication ends, these tubules morph into membranous assemblies that appear as vesicular-tubular clusters when cross-sections are imaged using electron microscopy (8, 90, 93). The enteroviral nonstructural protein 2(B)C is a member of the helicase superfamily 3 (94–96). This protein assembles into hexameric rings (96) and brings membranes together (90), potentially creating a channel through which the viral genome can be translocated from within the array of virus-induced membranes to the cytoplasm (90, 97). When processed capsid precursors accumulate to a sufficient level, functional intermediates form, perhaps half capsids (46), that may be localized to the 2(B)C channels by interactions of the capsid protein, VP3, with 2(B)C (98, 99). Empty capsids may then be filled with genomic RNA to produce virions. Virions are trafficked selectively into autophagosomes; empty capsids are excluded (21). Loading of virions into autophagosomes requires microtubule-associated protein 1B-light chain 3 (LC3B) and/or GABA Type A Receptor-Associated Protein (GABARAP) (18, 19, 21). The virion, 2(B)C, and 3CD can be pulled down in association with LC3 (24). Once in autophagosomes, cloaked virions traffic to the cell periphery and are ultimately released from cells in single-membrane vesicles after autophagosomes (or amphisome-like vesicles) fusion with the plasma membrane (19, 100).

Our data suggest that PV 3CD protein contributes to the last three steps described above. These steps are (1) movement of particles from the site of assembly into autophagosomes; (2) proper formation of autophagosomes; and (3) movement of autophagosomes containing PV virions from the perinuclear region of the cell to the periphery.

Using antibodies with some capacity to distinguish between empty capsids and virions, we were able to use immunofluorescence analysis of WT PV-infected cells to monitor spatiotemporal changes during morphogenesis (**Fig. 3**). By 4 hpi, empty particles had accumulated in the perinuclear region of the cell (WT 4 hpi in **Figs. 3C,D**), transitioning to virions by 6 hpi (WT 6 hpi in **Figs. 3C,D**). Once matured, virions moved from the perinuclear region of the cell to the periphery (WT 6 hpi in **Figs. 3C, D**). 3CD protein tracked with virus particles, moving from the perinuclear region of the cell to the periphery (WT 4 hpi and 6 hpi in **Figs. 3C, D**). We did not observe 3AB protein movement (data not shown). By 8 hpi, levels of virions and 3CD diminished substantially, consistent with both being released from the cell (WT 8 hpi in **Figs. 3C, D**). Although particle maturation occurred normally for F30Y PV, at least as measured by immunofluorescence, virions, and 3CD were trapped in the perinuclear region of the cell (**Figs. 4A, B**). We suggest the existence of a physical interaction between 3CD and virions that facilitates virion incorporation into vesicular carriers used for non-lytic spread.

Both WT and F30Y PVs appeared identical in their ability to induce autophagic signals based on LC3 lipidation and cleavage of an LC3 adaptor protein (**Figs. 5E, F**). For WT PV, we observed a clear colocalization of LC3 with virions (**Figs. 5G, H**) and 3CD protein, based on the association of virions and 3CD shown above (**Figs. 3C, D**). This colocalization was maintained from the perinuclear region at 4 hpi (**Figs. 5G, H**) to the periphery at 6 hpi (**Figs. 5G, H**). The association was no longer detectable, most likely because of the release of vesicular carriers of the virus that presumably also contained 3CD. For F30Y PV, colocalization of LC3 with virions and 3CD was lost (**Figs. 5I, J**).

These data are consistent with our proposition that an interaction between virions and 3CD delivers virions into LC3-marked vesicles. The 3CD derivative cannot facilitate virion loading. Interestingly, GABARAP-mediated loading of virions appeared unaffected for F30Y PV, suggesting the existence of two independent mechanisms for non-lytic spread (**Figs. 6C, D**). Two mechanisms would also explain our observation that not all non-lytic spread was eliminated for F30Y PV (**Fig. 2H**).

We have used high-angle annular dark-field (HAADF) scanning transmission electron microscopy (STEM) on a 200 kV instrument for the first time to monitor PV-infected cells which may represent its first use in characterizing virus-infected cells. HAADF STEM permitted us to visualize all of the intermediates and products expected for a virus using secretory autophagy for non-lytic spread (**Fig. 7**). Virion formation was not essential for formation of autophagosomes as they formed in the presence of hydantoin (**Fig. 8A**).

However, the 3CD derivative prevented formation of membranous structures expected if the sole defect were related to virion loading (**Fig. 8C**). We suggest a role for 3CD in virus-induced autophagosome biogenesis independent of its role in virion loading. LC3-interacting regions (LIRs) are predicted in all four capsid protomers (**Fig. S2**) and 3CD (**Fig. 9A**). Capsid, 2(B)C and 3CD proteins coprecipitate with LC3 (24).

Interestingly, there is an LC3 epitope located at the cleavage site between VP4 and VP2 of the VP0 capsid precursor that interacts with the LC3 adaptor protein SQSQTM1/p62 and may be capable of interacting with other LIR-containing proteins (89). Viral LIRs may mediate capsid-LC3 or capsid-3CD-LIR interactions required for virion loading.

F30Y increased the affinity of this highly conserved LIR (**Fig. 9H**) and perturbed the local conformations of both the 28-33 and 216-221 LIRs (**Fig. 10**). Such conformational changes may interfere with 3CD-dependent loading. These conformational changes may also interfere with autophagosome formation in the absence of cargo loading by preventing cargo-independent interactions of 3CD with LC3.

If one listed the function of 3CD protein 20 years ago, at the top of the list, functions related to its protease activity or its viral RNA-binding properties would appear (101). Over the past two decades, it has become clear that 3CD has many functions before genome replication related to its phospholipid-binding activity (8, 9, 102). These include the ability to induce phosphatidylinositol-4-phosphate and phosphatidylcholine synthesis and membrane biogenesis, which are presumably required to form the organelles for genome replication and virus assembly (8, 9, 91). Here, we have uncovered 3CD contributions after genome replication. The determinants of 3CD underpinning these functions need to be clarified but may include its LIRs. A much more deliberate analysis of LIRs encoded by enteroviruses is warranted. The ability of 3CD to exhibit so many diverse functions is likely related to the extraordinary conformational dynamics of this protein (26, 28, 84) and the sensitivity of these dynamics to even single amino acid substitutions, as shown here (**Fig. 10**). Together, this study highlights a crucial example of how LIR and LIR-like sequences in the viral genome play a significant role in hijacking components of the autophagy pathway to multiply and spread and underscores the importance of further exploring these sequences in other viral proteins and their potential function in a multitude of viruses with similar mechanisms.

## MATERIALS AND METHODS

### Cells and cultures

HeLa cells (CRM-CCL-2) were purchased from the American Type Culture Collection (ATCC) and grown in Dulbecco’s Modified Eagle Medium: Ham’s nutrient mixture F-12 (DMEM:F12) (Gibco). HAP1 human near-haploid cells were purchased from Horizon Discovery Group (Horizon) and grown in Iscove’s Modified Dulbecco’s Medium (IMDM). All cell lines were supplemented with 10% heat-inactivated fetal bovine serum (HI-FBS) (Atlanta Biologics), 1% of a penicillin and streptomycin mixture (P/S) (Corning), and maintained at 37° C and 5% CO_2_.

### Viruses

Wild-type and recombinant viruses were recovered following in vitro transcribed RNA transfection in HeLa cells. RNA was produced from full-length cDNA as described in the "Plasmids, in vitro transcription, cell transfection, and virus quantification" section. PV type 1 (Mahoney) was used as our WT PV strain throughout this study. The virus was quantified by standard plaque assay methods yielding virus titers (pfu/mL).

### Antibodies

The following commercially available and in-house produced antibodies were used at the specified dilutions in this study: human monoclonal A12 (gift from the Altan-Bonnet and Amy Rosenfeld Labs) (1:10,000 or 1:1500), mouse monoclonal Mab234 (gift from Andrew McAdam) (1:800), rat polyclonal PV 3CD (1:800), rabbit polyclonal PV 3AB (1:800), rabbit polyclonal PV VP2 (Cameron) (1:1000), rabbit monoclonal LC3B-D11 (Cell Signaling) (1:200), mouse monoclonal LC3B (Cell Signaling) (1:100), GABARAP monoclonal (Cell Signaling) (1:100), GABARAP monoclonal (ProteinTech), SQSTM1/p62 rabbit (Cell Signaling) (1:1000), rabbit αTubulin (Cell Signaling) (1:1000). Antibodies against PV 3D and 3AB were produced in Cameron Lab. All secondary antibodies goat anti-(human, mouse, rat, or rabbit) (H+L) used for immunofluorescence (1:1000) were purchased from Invitrogen. Secondary antibodies for western blotting: rabbit anti-HRP (Amersham GE Healthcare) and mouse anti-HRP (Cell Signaling).

### Reagents

Where specified, guanidine hydrochloride (GuHCl) (Sigma) was added to growth medium at 3 mM to inhibit PV genome replication, and 5-(3,4-dichlorophenyl) methylhydantoin (hydantoin) (Enamine) at 50 µg/mL to inhibit post-replication steps of PV infection.

### Plasmids, in vitro transcription, cell transfection, and virus quantification

Subgenomic WT and replication-incompetent GAA PV replicons were previously described (37). All insertions/deletions were produced using overlap extension PCR or gBlock gene fragments from Integrated DNA Technologies (IDT). Desired insertions/deletions or mutations in the PV cDNA were verified by DNA sequencing. For PV unaG_pv,_ the unaG-coding sequence was embedded between the 2C/3A coding region. The unaG-encoding sequence contained a 3C protease cleavage site at its carboxyl terminus for proteolytic cleavage/release of unaG protein engaged by 3C protease activity. Plasmids encoding PV genomes (full-length or subgenomic) were linearized using an *ApaI* restriction enzyme site.

All linearized cDNAs were *in vitro* transcribed using a T7 RNA polymerase produced in Cameron Lab and treated with 2 units of DNAse Turbo (ThermoFisher) to remove the residual DNA template. The RNA transcripts were purified using RNeasy Mini Kit (Qiagen) before spectrophotometric quantification. Purified RNA in RNase-free H_2_O was transfected into cells by electroporation using a Bio-Rad instrument (Gene Pulser).

Virus yield was quantified in HeLa cells by plaque assay. Cells and/or supernatant media were harvested post-transfection or infection at the specified time points, subjected to three freeze-thaw cycles, and clarified by ultracentrifugation. The supernatant was seeded on a fresh HeLa cell monolayer in 6-well plates and incubated at room temperature for 30 min before rinsing with 1X PBS. Then, a 1% (w/v) low-melting agarose/media overlay was added. Cells were incubated for either 2 days using WT PV or 3 days using F30Y PV, then fixed and stained using a PFA-containing crystal violet solution. Plaques were quantified to yield a PFU/mL titer.

### PV RdRp Biochemical characterization

Reactions were performed essentially as described in (29). All reactions contained 25 mM HEPES pH 7.5, 5 mM MgCl2, 10 mM BME, 60 µM ZnCl2, and 50 mM NaCl.

Reactions were performed at 30 °C. Reactions were quenched by adding EDTA to a final concentration of 50 mM. An equal volume of loading buffer (90% formamide, 0.025% bromphenol blue, and 0.025% xylene cyanol) was added to quenched reactions, and products were resolved from substrates by denaturing PAGE and visualized by using a PhosphorImager (GE) and quantified by using ImageQuant TL software (GE). For all reactions, the formation of 11-mer product RNA was monitored. Kinetics of complex assembly: Reactions contained 2 μM primed-template RNA substrate S/S-U (1 μM duplex), 500 μM ATP, and 1 μM PV RdRp. Reactions were initiated by adding PV RdRp and quenched at various times. Active site titration: Reactions contained 20 μM primed-template RNA S/S-U (10 μM duplex), 500 μM ATP, and 2 μM PV RdRp. Reactions were initiated by adding PV RdRp and quenched at various times. Kinetics of complex dissociation: 2 μM PV RdRp was incubated with 5’-P-labeled S/S-U (1 μM duplex) for 90 sec to assemble enzyme-RNA complex, then, unlabeled S/S-U (trap) was added to a final concentration of 100 μM. At various times after the addition of trap RNA, the amount of complex remaining was determined by taking a reaction aliquot and rapidly mixing it with an equal amount of 1 mM ATP. After mixing with ATP, the reactions were allowed for 30 sec and quenched.

### Sub-genomic replicon luciferase assay

Subgenomic replicon luciferase assays were performed as described previously (8). Subgenomic replicon RNA (5 μg of in vitro transcribed RNA) was electroporated into HeLa cells. The cells were incubated in standard growth media (DMEM/F12 supplemented with 10% fetal bovine serum and 1% penicillin/streptomycin; cells were harvested and lysed using 100 μL of 1X cell culture lysis reagent (CCLR, Promega) at the indicated times post-electroporation. Luciferase was measured as a surrogate for genome replication using a relative light unit (RLU) normalized to protein content (µg) from an absorbance measure of the collected lysates. Luciferase activity was measured by adding an equal volume of firefly luciferase assay substrate (Promega) to cell lysates and measured in a Junior LB 9509 luminometer (Berthold Technologies) or a BioTek plate reader.

### Plaque assay comparing plaque forming unit phenotypes

HeLa cell monolayers were infected with 50 plaque-forming units (PFU) using WT or F30Y PV. Cells were then incubated at 37 °C for three days ahead of staining with crystal violet for plaque quantification and phenotype assessment.

### One-step growth curve of media-associated and cell-associated viruses

HeLa cell monolayers were infected with WT or F30Y PV at an MOI of 10. The virus was then collected from the supernatant and cells (independently) at the indicated time points. Media-associated (supernatant) and cell-associated (cells) virus titer was determined by plaque assay.

### Cell-free PV synthesis supplemented with purified 3CD protein

Cell-free PV synthesis experiments were carried out as described by Franco et al. (2005). HeLa cytoplasmic extracts (cell-free) were supplemented with viral RNA as a translation template in the presence of unlabeled methionine, 200 μM each CTP, GTP, UTP, and 1 mM ATP. Exogenous purified WT or F30Y PV 3CD protein was introduced to the reaction. After a 12–15 hr incubation, samples were diluted with phosphate-buffered saline and applied to HeLa cell monolayers. Virus titers were determined by plaque assay.

### Bulk spread assays

HeLa cells in suspension were stained using a membrane dye Vybrant DiD (Molecular Probes) and infected with a green fluorescence PVeGFP_pv_ reporter variant at an MOI of 5. Infected/dyed cells (red) were seeded on top of a naïve HeLa cell monolayer. Fluorescence is monitored over time to detect both primary and secondary infections. Primary infected cells were observed and depicted in yellow when green (eGFP expression) and red signal (cell dye) colocalized in overlays. Spread was detected when a secondary wave of PV green fluorescence signal (green only) originating from the newly infected monolayer of unstained cells was observed.

For imaging, the plate was placed in the chamber of a WSKM GM2000 incubation system (Tokai, Japan), which was adapted to a Nikon Eclipse Ti inverted microscope (Nikon, Japan). Automatic bright-field and fluorescence imaging were performed every 30 minutes from 3 to 24 hpi with a ProScan II motorized flat top stage (Prior Scientific, USA), a CFI60 Plan Apochromat Lambda 10× objective, and a Hamamatsu C11440 camera. Image analysis was performed using the ComDet module. Cells were quantified as GFP or Red Dye positive given a fluorescence intensity threshold; the number of green cells was normalized to the initial fraction of GFP-positive cells and plotted.

### Immunofluorescence assays

HeLa cells were grown in coverslips, treated as described in the respective figures, and fixed at the specified time points using 4% formaldehyde in PBS for 20 min. Immunostaining was performed by permeabilizing with 0.2% Triton X for 10 min, blocking with 3% Goat Serum in PBS for 1 hour, and incubating in primary antibodies for 1 hour. Following washes, cells were incubated with secondary antibodies for 1 hour and either DAPI (Sigma) or TOPO-3 (Invitrogen) for 10 mins. The processed coverslips were mounted on glass slides using ProLong Glass Antifade Mountant (Thermo Scientific).

Imaging was performed using an oil immersion 63X objective on the Zeiss 880 confocal microscope at the Hooker Imaging Core at UNC. Images were acquired and minimally processed using the Zeiss Zen software. Multiple images were obtained, and a representative cell was selected from representative image fields.

### Fluorescence intensity profiles

A white line extending from the nuclear envelope to the plasma membrane of cells was drawn for "profile fluorescence" signal quantification. Intensity profile measurements were taken from regularly spaced points along a line segment to depict the spatial and temporal dynamics of fluorescence reactivity, levels, and signal overlap in infected cells over time, using the "Profile" module in the Zeiss Zen software. Values were plotted as a smooth line graph with relative fluorescence intensity units (RFU) on the Y-axis, and the distance (nm) of each fluorescence signal was plotted as independent lines in the graph. 3 to 5 separate cells in each representative image field were quantified to determine the intensity profile pattern of each collected condition and time point.

### Single-cell spread assays

Cells in suspension infected with a reporter PV-unaG_pv_ virus variant (green). Infected cells were paired with stained (Vybrant DiD) uninfected cells (red) in isolated chambers of a multi-chamber microfluidics polydimethylsiloxane (PDMS) device as described (59, 103). In this study, this device was modified to harbor cell pairs (103). Fluorescence is monitored over time to detect an initial wave of infected cells expressing green fluorescence., yielding a yellow fluorescence overlay (see yellow cells). Spread was detected when a secondary wave of green fluorescence signal was observed in red-dyed cells, producing a colocalized yellow signal. Spread events were further extrapolated into no-spread, lytic spread, and non-lytic spread. In no spread, no secondary infection signal was detected after a primary cell green fluorescence signal. In lytic spread, the secondary infection signal arose after losing the primary cell green fluorescence (lysis). In non-lytic spread, the secondary infection signal was detected while green fluorescence was still present in the primary infected cell. HeLa or HAP1 cells were infected with either WT or F30Y PVunaG_pv_ at an MOI of 5 and paired with uninfected stained cells (red).

For imaging, the microfluidics device was placed in the chamber of a WSKM GM2000 incubation system (Tokai, Japan), adapted to a Nikon Eclipse Ti inverted microscope (Nikon, Japan). Automatic bright-field and fluorescence imaging were performed every 30 minutes from 3 to 24 hpi with a ProScan II motorized flat top stage (Prior Scientific, USA), a CFI60 Plan Apochromat Lambda 10× objective, and a Hamamatsu C11440 camera. The fluorescence intensity of single cells and the background intensity of the microwells were extracted with a customized MATLAB script. Relative intensity was calculated as (Cell intensity - Background)/Background. No spread, lytic, and non-lytic events were quantified as percentages of the total events. The values were represented as mean ± standard error (SEM) from an n=3. Significant differences between conditions were noted based on a student’s t-test with p-values below 0.05.

### Immunoblotting

Cells were lysed in radioimmunoprecipitation assay (RIPA) buffer containing an inhibitor cocktail of phenylmethylsulfonyl fluoride (PMSF) (1:100) (American Bioanalytical), Protease Inhibitor Cocktail (Sigma-Aldrich) (1:100), and Phosphatase Inhibitor Cocktail I (Abcam) (1:100). Lysates were collected and clarified by centrifugation. The lysate was mixed with 4× Laemmli buffer (Bio-Rad), boiled, and processed by SDS-PAGE. The samples were then transferred from the gel to a 20 μm nitrocellulose membrane (Bio-Rad) using the TurboBlot system (Bio-Rad). Membranes were blocked in Everyday Blocking Reagent (Bio-Rad) and probed with anti-LC3B (1:1000), anti-SQSTM1/p62 (1:1000), anti-VP2 (1:5000), or anti-tubulin (1:5000) antibodies overnight. Anti-rabbit or mouse-HRP was used as a secondary antibody at a 1:5000 dilution. Protein bands were visualized with the ECL detection system (Bio-Rad) using the ChemiDoc MP imaging system (Bio-Rad). Basic post-imaging editing and band quantification were performed using the Bio-Rad Image Lab software.

### Scanning Transmission Electron Microscopy (STEM)

HeLa cells were infected, fixed, and embedded for TEM studies, as described previously (41). Briefly, cells were harvested and fixed with 1% glutaraldehyde, washed with 0.1 M cacodylate (sodium dimethyl arsenate, Electron Microscopy Sciences) twice for 5 min each, incubated in 1% reduced osmium tetroxide containing 1% potassium ferricyanide in 0.1 M cacodylate for 60 min in the dark with one exchange and washed two times with 0.1 M cacodylate again. *En bloc* staining was performed with 3% uranyl acetate in 50% ethanol for 60 min in the dark. Dehydration was carried out with varying ethanol concentrations (50, 70, 95, and 100% for 5–10 min) and 100% acetonitrile.

Embedding was performed overnight with 100% Epon at 65° C. The embedded sample was sectioned with a diamond knife (DiATOME) to slice it to a 60–90 nm thickness using an ultramicrotome (Reichart-Jung). The sectioned sample was placed on a copper grid (Electron Microscopy Sciences) and stained with 2% uranyl acetate in 50% ethanol, followed by lead citrate staining for 12 min. The grid was washed with water and dried thoroughly.

HAADF-STEM (High Angle Annular Dark Field - Scanning Transmission Electron Microscopy) images were collected using a Thermo Scientific Talos F200X G2 (S)TEM operated at 200 kV and a beam current of approximately 0.12 nA. The Talos (S)TEM instrument has a resolution limit of 0.16nm in STEM mode, providing enhanced contrast compared to TEM (roughly proportional to Z^2). The contrast is also reversed when compared to TEM, with the vacuum appearing dark. Before STEM imaging, the grid square was first "beam showered" in TEM mode at a maximum beam current for approximately 10 minutes, with the beam spread to cover one entire grid square. This reduces the carbon contamination build-up that is naturally present on the surface of all samples (104). By beam showering, we lessen the contamination build-up that would otherwise limit contrast in the STEM image.

### LC3-interacting region (LIR) predictions

LIR predictions were carried out as described by Jacomin et al., using the iLIR database (https://ilir.warwick.ac.uk) developed by the Nezis group at Warwick U.K (81). The consensus sequences of selected enteroviruses used in this analysis included: PV (Genbank:_V01149.1), CV-A16 (GenBank:_U05876.l), EV-71 (GenBank:_U22521.1), CV-B3 (GenBank:_M88483.l), EV-D68 (GenBank:_AY426531.l), EV-E1 (GenBank:_D00214.1), EV-F1 (GenBank:_DQ092770.l), EV-G1 (GenBank:_AF363453.l), EV-H1 (GenBank:_AF201894.l), EV-J GenBank:_AF326766.2), HRV-A1 (GenBank:_FJ445111.l), HRV-B3 (GenBank:_DQ473485.l), HRV-C1 (GenBank:_EF077279.1), and HRV-B14 (GenBank:_U05876.l).

### Protein structure analysis

PV 3C and 3D LC3 Interaction Regions (LIRs) were computationally scrutinized using AI and the Alphafold multimer server (82). The exposed consensus motif (W/F/Y) (x) (x) (L/I/V) within the LC3-interacting regions of 3C and 3D, as from their crystal structures, underwent structural analysis using PyMOL software (The PyMOL Molecular Graphics System, Version 2.0 Schrödinger, LLC). Thirteen LIRs within the 3CD protein were identified, and their sequences were verified for conservation through a Blast alignment of all Enteroviral sequences. Notably, two LIRs in the palm and thumb domain of the 3D structure were strictly conserved across multiple Enterovirus species.

### Alphafold docking and binding

Complexes of PV 3D and LC3A/LC3B/Gabarap were made using the Alphafold multimer docking server, incorporating the sequences of 3D and LC3A/LC3B/Gabarap. Similar complexes were predicted for PV 3C and LC3A/LC3B/Gabarap. The five complex models generated by Alphafold underwent analysis for their scores and interface region, with the top model being energy minimized using the Yasara energy minimization server (83).

As from the best binding complexes identified, specific PV 3D LIR peptides in regions 30-34, namely LEPSAF(30)HYVFEGVK and F30Y variant LEPSAY(30)HYVFEGVK, along with LIR region 218-221 CDPDLFWSKIPVLM, were assessed for their binding affinities with LC3B. Peptide and LC3B Alphafold-multimer docking models were created, and the best model of the complex underwent energy minimization in Yasara. The computed DeltaG and Kd for the resulting complexes were deduced using the Prodigy software (105).

### Molecular Dynamics Simulations

WT PV 3CD and F30Y PV 3CD MD simulations were performed using the AMBER software suite (106), applying parameters from amber forcefield 14SB (107). The 3CD WT monomer coordinates were extracted from the 3CD protein (PDB 2IJD) (39) crystal structure and prepared for simulations as described previously (28). The F30Y PV 3CD system was prepared in silico by replacing Phe at position-30 with Tyr; any steric clashes produced in the prepared mutant were removed by subsequent energy minimization and equilibration during MD simulations.

All-atom MD simulations were performed in explicit water (TIP3P model (108)); a minimal distance of 20 Å between the edge of the solvent box and any protein atoms was imposed. A cutoff radius of 12 Å was used to calculate non-bonded interactions with periodic boundary conditions applied; the particle mesh Ewald method (109, 110) was used to treat electrostatic interactions. The SHAKE algorithm (111) was employed to constrain hydrogens bonded to heavy atoms. The simulations were performed by first relaxing the systems in two cycles of energy minimization; subsequently, the systems were slowly heated to 300 K using the parallel version PMEMD under NVT conditions (constant volume and temperature). Langevin dynamics (112) with collision frequency (γ=2) were employed to regulate temperatures. The heated systems were then subjected to equilibration by running 100 ps of MD simulations under NPT conditions (constant pressure and temperature) with 1 fs integration time step. MD trajectories were collected over 200 ns at 1 ps interval and 2 fs integration time step. Analyses of the trajectories from MD simulations were done using the CPPTRAJ program (113). MD simulations were carried out on a multi-GPU workstation with 2x AMD EPYC 7702 64-core processor and 2x Nvidia RTX A5000.

## ACKNOWLEDGMENTS

This work was supported by funding from the NIH NIAID R37AI053531, F31AI179022, and Burrough’s Wellcome Fund BWF1022057. Microscopy was performed at the UNC Hooker Imaging Core Facility, supported in part by P30 CA016086 Cancer Center Core Support Grant to the UNC Lineberger Comprehensive Cancer Center—special thanks to Director Wendy Salmon and EM Research Specialist Paul Risteff. The co-authors acknowledge the use of the Penn State Materials Characterization Lab and Huck Institutes of the Life Sciences Microscopy Core Facility. Special thanks to Director Grang Ning and EM sample specialist Missy Hazen.

## SUPPLEMENTAL MATERIAL

Movie S1

Movie S2

---

**Figure S1.**
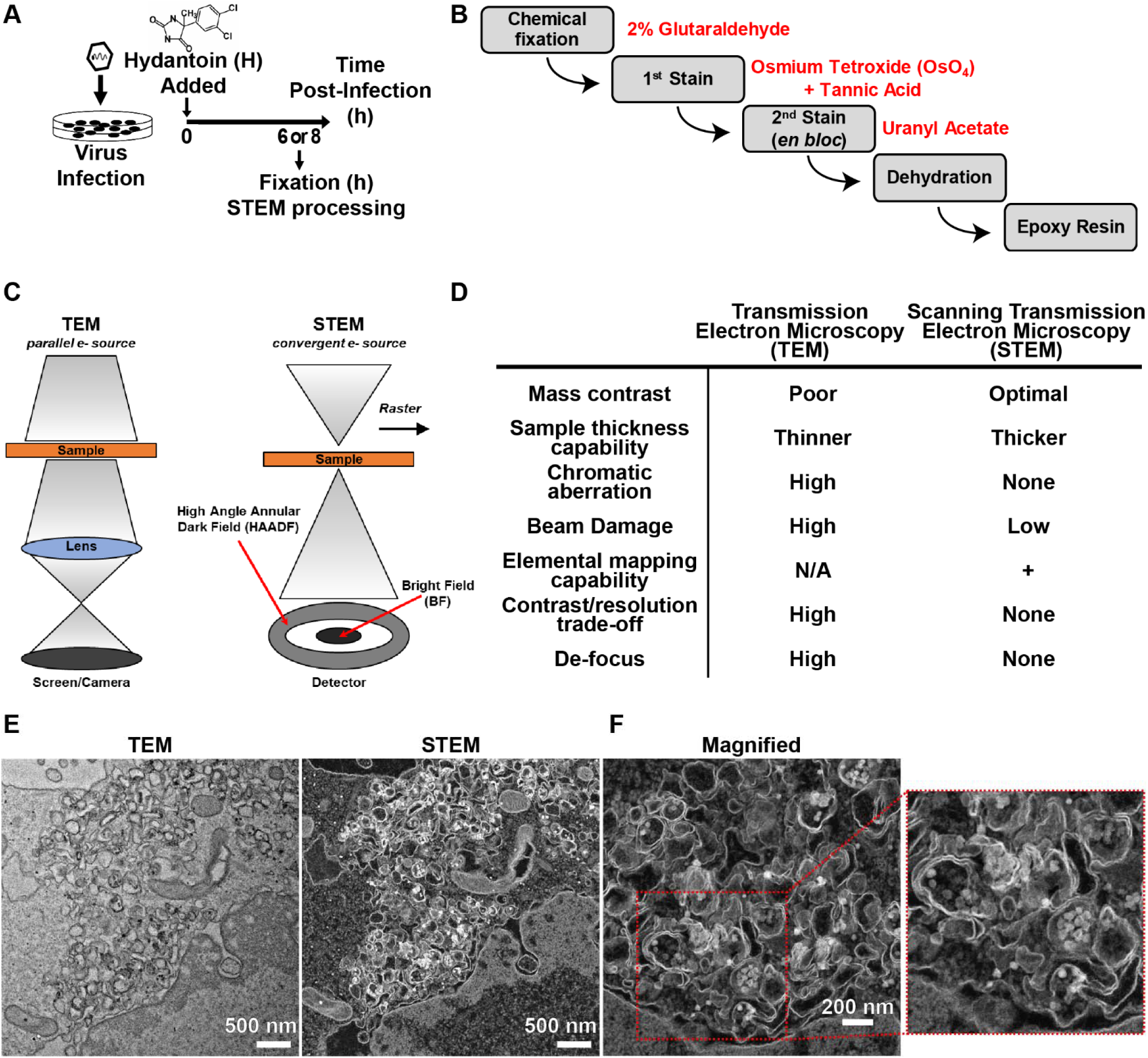
An alternate imaging approach: Scanning transmission electron microscopy (STEM). **(A) Cell lysate preparation for STEM.** Infection of HeLa cell monolayers was carried out in the presence or absence of hydantoin. A cell suspension is then prepared using trypsin to release the monolayer at the stated time points 6 or 8-hours post-infection. Cells are gently pelleted, fixed, and processed as described in panel **(B)**. **(B) Cell microsection preparation for STEM.** Cell pellets were subjected to chemical fixation using 2% glutaraldehyde. An initial stain was performed using osmium tetroxide, followed by tannic acid treatment. A second *en bloc* stain was completed using uranyl acetate. Cell pellets were then dehydrated and embedded in an epoxy resin. Thin microsections were then collected and placed on a carbon-coated grid, where a third and final on-grid stain was performed. **(C) Schematic of TEM and STEM microscopy**. TEM is set up much like light microscopy but uses electrons and electromagnetic lenses instead of light. Briefly, the beam hits the sample, electrons are scattered, and the lens forms an image projected to the camera. STEM is entirely different. The beam is converged to a single point, then rastered across the sample, and a detector collects the resulting scattered electrons. In short, TEM contrast comes from unscattered electrons. In STEM, contrast comes from scattered electrons. **(D) Advantages and disadvantages of TEM and STEM imaging.** This table discusses the advantages and disadvantages of Transmission Electron Microscopy (TEM) and Scanning Transmission Electron Microscopy (STEM) to provide some perspective on the factors influencing the contrast gains obtained when imaging biological samples using STEM. In short, we enumerate several advantages of using STEM imaging in the ultrastructural analysis of biological samples, such as membrane derangements in infected cells. **(E) TEM and STEM imaging mode comparison.** HAADF-STEM (High Angle Annular Dark Field - Scanning Transmission Electron Microscopy) imaging of WT PV-infected HeLa cells. HeLa cells were infected with WT PV at an MOI of 10 and then fixed in glutaraldehyde 6 hours post-infection (hpi). Fixed samples were dehydrated, stained, embedded, and sectioned in thin micrographs for imaging as described in panels **(A)** and **(B)**. Images were collected using a Thermo Scientific Talos F200X G2 (S)TEM operated at 200 kV and a beam current of approximately 0.12 nA. The contrast is also reversed when compared to TEM, with the vacuum appearing dark. WT infection induces virus-containing double membranous vesicles and multi-vesicular amphisome-like vesicles with virions in the intra-luminal vesicles. Large outer vesicles with intra-luminal vesicles (100-300 nm diameter) contain ∼30 nm particles inside. Double membrane vesicles are located at sites where vesicular-tubular clusters are observed in TEM mode. **(F) STEM imaging of WT PV-infected HeLa cells (magnified)**. In this magnified view, we look closely at observed structures in panel **(E)**. Large outer vesicles with intra-luminal vesicles (100-300 nm diameter) contain ∼30 nm particles inside. Double membrane vesicles are located at sites where vesicular-tubular clusters are observed in TEM mode. 30 nm virus particles observed inside of intra-luminal vesicles. Close-up view of an intra-luminal vesicle that contains 30 nm particles.

**Figure S2.**
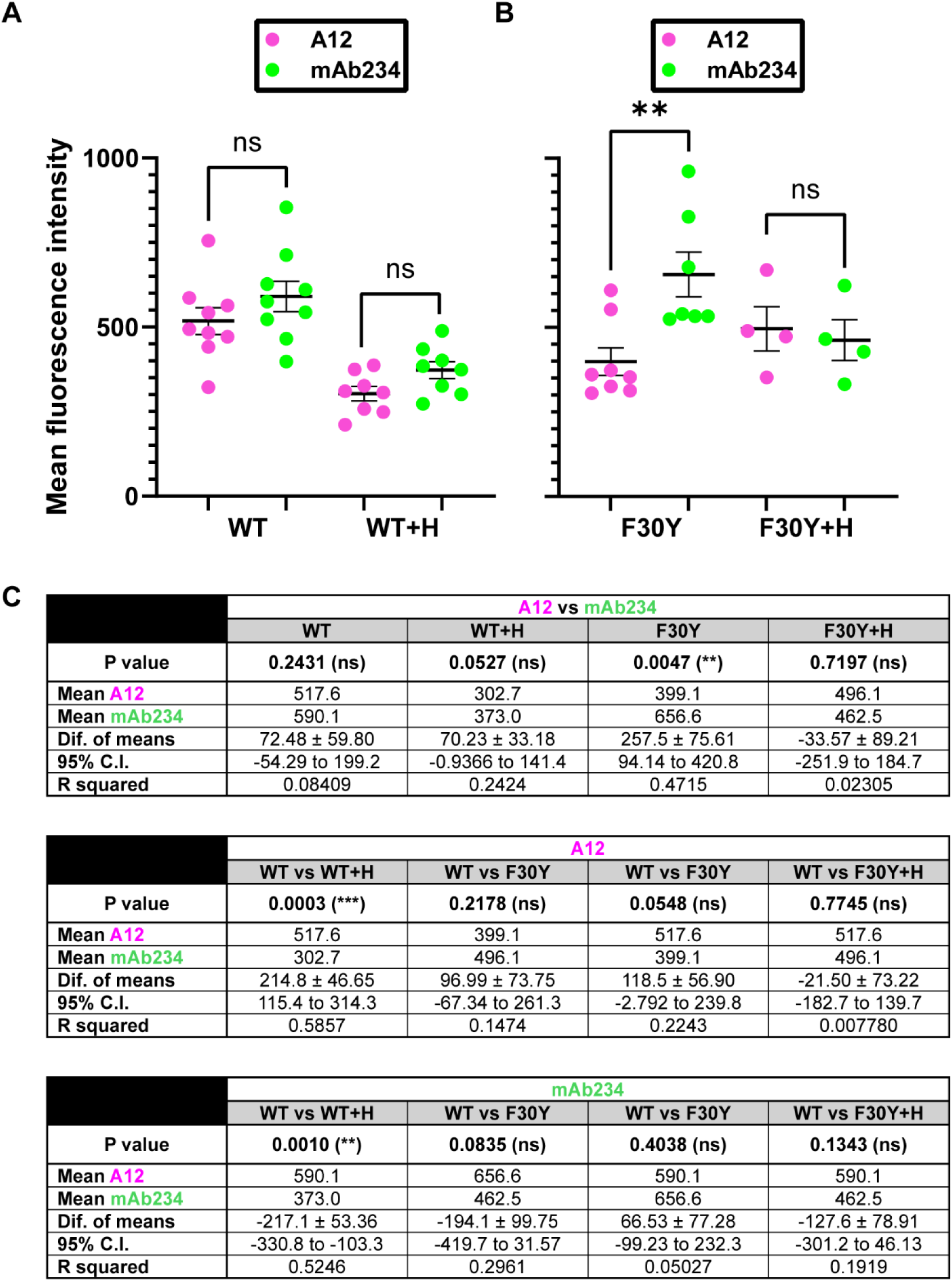
A12 and mAb234 fluorescence intensity analysis of PV-infected cells. **(A) Confocal immunofluorescence imaging intensity measurements of A12 and MAb234 in WT PV-infected HeLa cells.** Images illustrate intensity measurements of whole cells in representative immunofluorescence image fields of WT-infected HeLa cells (MOI of 10) in the presence and absence of hydantoin, as described in **Fig 2C**. Mean fluorescence intensity is plotted on the y-axis, and the conditions on the x-axis. **(B) Confocal immunofluorescence imaging intensity measurements of A12 and MAb234 in F30Y PV-infected HeLa cells.** Images illustrate intensity measurements of whole cells in representative immunofluorescence image fields of F30Y-infected HeLa cells (MOI of 10) in the presence and absence of hydantoin, as described in **Fig 2C**. Mean fluorescence intensity is plotted on the y-axis, and the conditions on the x-axis. **(C) Statistical analysis on fluorescence intensity measurements.** An unpaired student t-test analysis was performed to compare the intensity measurements of WT and F30Y-infected cells in the described conditions. A p-value lower than 0.005 was considered significant with a 95% confidence interval.

